# Baseline regulatory programs in larval and adult neural progenitors converge towards an injury-induced state after spinal cord injury

**DOI:** 10.64898/2026.07.24.740552

**Authors:** Mehmet Ilyas Cosacak, Dmitrii Severinov, Markus Westphal, Kim Heilemann, Denise Bärhold, Anja Bretschneider, Susanne Reinhardt, Juliana G. Roscito, Thomas Becker, Catherina G. Becker, Anna R. Poetsch

**Author notes:** Contributed equally (Mehmet Ilyas Cosacak, Dmitrii Severinov).

## Abstract

Regeneration after spinal cord injury requires progenitor cells to convert injury-associated signals into coordinated remodeling of gene regulatory programs. Mammalian spinal progenitors show limited neurogenic output after injury, whereas zebrafish regenerate spinal neurons and recover motor function. To investigate the regulatory changes that allow ependymo-radial glia (ERG) cells, the progenitor cells of the zebrafish spinal cord, to generate new neurons, we combined single-nucleus gene expression and chromatin accessibility profiling across embryonic, larval, and adult stages with topic-based gene regulatory network (GRN) inference. We found that larval and adult ERGs enter the injury response from distinct regulatory baselines: larval progenitors are characterized by a gliogenic program, whereas adult progenitors maintain a comparatively quiescent state. Following injury, both populations gradually change their baseline programs and shift towards a lesion-associated module marked by stress-responsive and chromatin-associated regulators, including *jun*, *hmga1a*, *hmga2*, *ybx1*, and *foxj1a*. The shift away from homeostatic states is supported by decreased expression of the Notch-associated regulators nuclear factor I A (*nfia*) and *hey1* in larvae, while in adults, downregulation of the same nuclear factor and other TFs such as *bhlhe41* is associated with quiescence exit. Pathway analysis showed stage-specific alterations after injury, characterized predominantly by extracellular signaling and cytoskeletal reorganization in larvae and by metabolic and translational remodeling in adults. Despite divergence from the homeostatic states, injury-induced larval and adult GRNs remain distinct from embryonic hERG regulatory programs. Thus, larval and adult progenitors follow different trajectories from their baselines towards a related lesion-reactive state, in which shared regeneration-associated features are acquired within respective contexts.

## Introduction

Regeneration of neurons after spinal cord injury requires progenitor cells to convert injury-associated signals into coordinated remodeling of gene regulatory programs. This process depends not only on which genes are expressed after injury, but also on how progenitor regulatory programs are reorganized in the injury context. In mammals, endogenous spinal progenitors proliferate after injury but show limited, if any, neurogenic output and predominantly contribute to scar-associated glial responses (*1*, *2*). By contrast, zebrafish regenerate neurons in both larval and adult stages and recover motor function after spinal cord injury. This process involves re-initiation of neurogenesis from the spinal cord progenitor cells, the ependymo-radial glial cells (ERGs), glial bridging, and maturation of newborn neurons (*3–7*). However, the developmental stages differ in baseline tissue composition, developmental maturity, immune environment, progenitor plasticity and regeneration timescale: larval injury occurs in still-developing and low-level neurogenic tissue, whereas the adult spinal cord is largely quiescent under homeostasis and takes longer to regenerate (*5*, *7*, *8*). These differences raise the question of whether larval and adult ERGs reach similar regeneration-associated cell states starting from distinct tissue contexts and how their regulatory programs are reorganized to allow such convergence.

Previous studies across regenerating systems have shown that regeneration can reuse developmental genes while also activating regeneration-specific programs (*9–11*). However, gene-centered views provide only a partial picture of this process. Cellular state transitions depend on combinations of transcriptional regulators (TRs) acting in parallel with chromatin states and *cis*-regulatory elements (CREs) to coordinate target gene activity. These interactions can be represented as gene regulatory networks (GRNs), which provide a framework to understand how cell states are established, maintained, and remodeled (*12–14*). In the context of regeneration, GRNs therefore help us to reconstruct and to study the regulatory architecture changes during a progenitor state transition.

Single-cell multiome sequencing enables simultaneous profiling of gene expression and chromatin accessibility from the same nucleus, allowing transcriptional changes to be connected to CREs. To investigate this regulatory reorganization, we generated a single-nucleus multiome (snRNA-seq and snATAC-seq) atlas of zebrafish spinal cord progenitor regeneration, covering embryonic, larval and adult stages, injury conditions, and time. We integrated TR expression, chromatin accessibility, and gene expression using single-cell Deep Multi-omic Regulatory Inference (scDoRI), a method for topic-based enhancer-linked GRN inference (*15*). scDoRI represents regulatory complexity as low-dimensional, interpretable features termed “Topics”, where each topic corresponds to a putative enhancer-mediated TR-gene network. This allowed us to model each cellular state as a mixture of network activities and to identify shared and state-specific gene regulatory modules beyond discrete cell-type categories.

Using this atlas, we examined how ERG cells transition from baseline to lesion-reactive programs in larvae and adults. We identified a shared injury-associated regulatory module that overlays distinct stage-specific baseline programs: a gliogenic program in larval progenitors and a quiescent program in adult progenitors. Our results suggest a model in which ERG activation is a continuous regulatory transition rather than a discrete switch. In this model, larval and adult progenitors converge on a related injury-induced state through different regulatory routes and the regulatory core of this state differs from embryonic hERGs.

## Results

### Joint snRNA-seq and snATAC-seq profiling reveals injury-induced cell states

To investigate the dynamic changes of gene regulatory programs upon injury in zebrafish spinal cord ERG cells, we performed a single-nucleus multiome assay (joint snRNA-seq and snATAC-seq in the same nucleus) using the Tg(her4.1)^y83Tg^ line (*16*) (**Figure 1a**). Embryonic samples were collected from whole trunks at 1 day post fertilization (dpf), whereas larval samples were prepared from the trunk tissue surrounding the lesion site after spinal cord transection (*5*, *17*). In adults, spinal cord tissue spanning three vertebrae rostral and caudal to the lesion site was dissected after injury (*18*). In total, we profiled 27 samples spanning three developmental stages, 10 time points and two conditions. To account for developmental changes at the larval stage, we collected age-matched control samples at 4, 5 and 7 dpf in addition to the 3 dpf control; for the 3 dpf + 6 hours post lesion (hpl) time point, the 3 dpf control was used, as limited developmental changes were expected within 6 hours. For adult samples, exact age-matched controls after lesion were not collected, because major transcriptional or epigenetic changes were not expected over these post-lesion intervals. After quality control of the single-nucleus data and doublet removal, 172,433 nuclei were retained for downstream analysis (**Supplementary Figure 1**).

**Figure 1.**
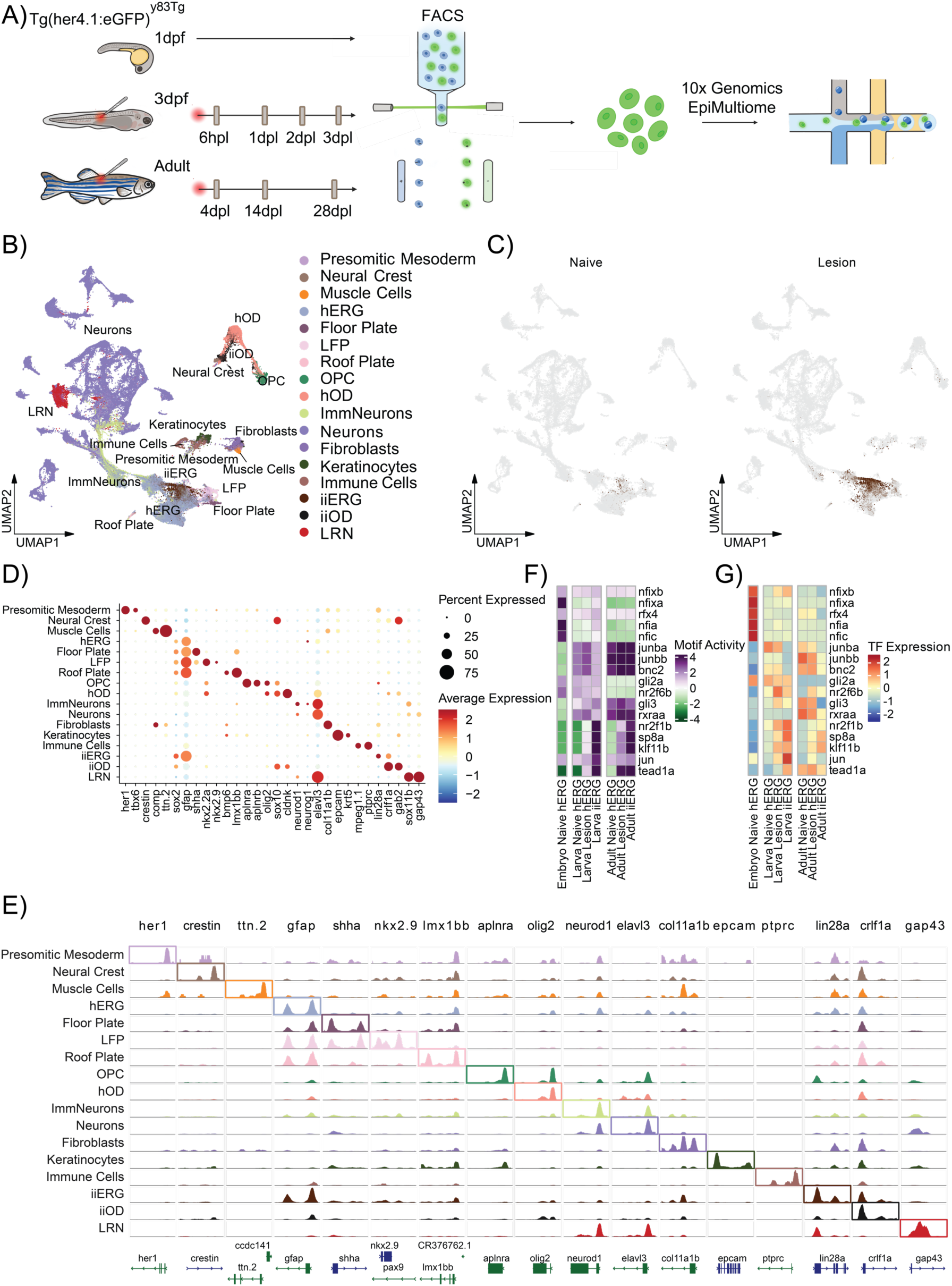
Single-nucleus multiome profiling resolves homeostatic and injury-induced cell states in the zebrafish spinal cord. (**A)** Experimental design for single-nucleus multiome profiling across developmental stages, injury conditions, and post-lesion time points. For each injury time point, control samples were collected. The 3 dpf samples served as controls for 3 dpf + 6 hpl, as developmental changes within 6 hours were considered negligible. Control samples at 4 dpf, 5 dpf, and 7 dpf were prepared for 3 dpf + 1 dpl, 3 dpf + 2 dpl, and 3 dpf + 4 dpl, respectively. Adult animals were expected to be transcriptionally and epigenetically similar; therefore, exact age-matched control samples were not collected for the Adult + 3 dpl, Adult + 14 dpl, and Adult + 28 dpl time points. dpf – days post fertilization, dpl – days post lesion, hpl – hours post lesion. (**B**) Integrated UMAP embedding of 172,433 nuclei annotated into 17 major cell populations. (**C**) UMAP embedding with highlighted 4391 iiERG cells across conditions; brown – iiERG cells, grey – the rest of the cells. (**D**) Average scaled gene expression of marker genes across annotated cell types. (**E**) Chromatin accessibility profiles at the TSS of selected marker genes. (**F**) Top five active motifs in progenitor cells at each developmental stage. The selection was done based on the average of the motif activity and corresponding TF expression. (**G**) Average scaled expression of selected TFs identified from motif activity analysis. hERG – homeostatic ependymo-radial glia; hOD – homeostatic oligodendrocytes; iiERG – injury-induced ERG; ImmNeurons – immature neurons; iiOD – injury-induced OD; LFP – lateral floor plate; LRN – lesion-reactive neurons; OPC – oligodendrocyte precursor cells.

To account for both stage-specific and shared cell states, we integrated the RNA and ATAC modalities for each developmental stage separately. Stage-specific annotations were then used in a joint atlas (**Supplementary Figure 2-6**). This strategy allowed us to directly compare related cell states while preserving the stage-dependent differences. Integration analysis revealed 17 major cell types (**Figure 1b**). ERG progenitors were annotated by expression of *gfap* and *sox2* (*19*) together with accessibility at the promoter regions of these genes (**Figure 1d,e**). In addition to homeostatic ERGs (hERG), we can detect a distinct injury-induced ERG (iiERG) population(*20*). The iiERG cluster is observed almost exclusively in lesioned samples and shows expression and accessibility of regenerative neurogenesis-associated genes *lin28a* (*20*, *21*) and *crlf1a* (*20*, *22*) (**Figure 1c-e**). In addition, Lateral Floor Plate (LFP) cells share several progenitor-associated features with ERGs. However, they form a distinct cluster and show enrichment of ventral floor plate markers: *nkx2.2a* and *nkx2.9* (*23–25*) (**Figure 1d**, **Supplementary Figure 2, 5**). Therefore, we annotated LFP separately and used ERG to refer to the progenitor cells outside the LFP cluster. We can also identify injury-induced oligodendrocytes (iiODs), defined by expression and accessibility of *crlf1a* and *gab2*. A lesion-reactive neuronal (LRN) population is enriched in larval injured samples and expresses the established neuronal regeneration-associated genes *sox11b* (*26*) and *gap43* (*27*) (**Supplementary Figure 4, 5**). We did not recover an adult LRN cluster, which likely reflects the cell capture strategy, as GFP-positive cells represent predominantly progenitor cells rather than their neuronal descendants. However, a transient population of injury-responsive neurons has recently been reported in the adult zebrafish spinal cord, especially one week post-injury(*28*, *29*). Overall, this atlas captures the target progenitor cell population in both homeostatic and injury-induced conditions across developmental stages, and in addition identifies oligodendrocyte and neuronal injury-associated states.

To define regulatory starting states from which ERGs respond to injury, we examined TF expression and motif activity across hERGs and iiERGs. Embryonic hERG cells show neural progenitor-like regulatory features, including activity of Gli family members and retinoid-associated factors (**Figure 1f,g**) (*30–33*). Larval and adult progenitors are characterized by the presence of nuclear factor I family members, including *nfia*, *nfixb*, *nfic*. However, these factors are associated with different developmental contexts: in larvae, *nfia* is linked to a gliogenic program, whereas *nfia* together with nfix genes was shown to be associated with quiescence in adults (*34–36*). In turn, *sp8a*, which is associated with pMN/p3 domain patterning (*37*), and context-dependent transcriptional activator or repressor *klf11b* (*38, 39*) show increased activity in larval hERGs (**Supplementary Figure 9b**).

Additionally, larval and adult hERGs differ at the transcriptional and epigenetic levels (**Supplementary Figure 9a,c,e**). The adult against larva comparison identified 3,057 differentially expressed genes (DEGs) in hERGs from uninjured animals and 2,392 DEGs in hERGs from lesioned animals. KEGG pathway analysis further highlighted the baseline differences in hERGs across stages (**Supplementary Figure 10**). Among hERGs from uninjured fish, adult cells show stronger enrichment for metabolic pathways, whereas larval cells are more strongly enriched for Notch- and translation-associated pathways. The comparison of hERGs from lesioned animals identified fewer significantly enriched pathways. Together with the lower number of DEGs, the reduced number of significant pathways might indicate a partial increase in transcriptional similarity between larval and adult hERGs after injury.

Despite these differences in baseline states, iiERGs from injured larvae and adults converge on a shared AP-1-associated response (*27*). Larval iiERGs additionally show a stage-specific increased expression level of *tead1a* (**Figure 1g**, **Supplementary Figure 9b**), indicating the regeneration-associated activity of Hippo signaling (*40*). At both developmental stages, reduced expression of nuclear factor I genes relative to hERGs indicates a shift away from the corresponding baseline regulatory programs. Furthermore, the adult against larva comparison of iiERGs identified approximately 35% fewer DEGs than the corresponding comparison of hERGs from uninjured animals and around 17% fewer DEGs than the comparison of hERGs from lesioned animals (**Supplementary Figure 9a**). At the pathway level, adult iiERGs show stronger enrichment for metabolic processes, whereas larval iiERGs are more enriched for signaling pathways, including Wnt, MAPK, and Cadherin signaling (**Supplementary Figure 10**). In general, these functional trends follow the stage-dependent patterns observed in hERGs, even though the specific enriched pathways slightly differ.

Notably, both motif activity and expression of TFs associated with embryonic hERGs are lower in iiERGs. Direct comparison of iiERG cells with embryonic hERGs revealed 3,892 DEGs and 19,874 differentially accessible regions (DARs) for larvae and 3,676 DEGs and 21,105 DARs for adults (**Supplementary Figure 9a,e**). Pathway enrichment analysis further supported both the divergence of iiERGs from embryonic hERG cells and the convergence of injury-induced state across developmental stages, as the pathways enriched or depleted in larval and adult iiERGs relative to embryonic hERGs show substantial overlap (**Supplementary Figure 10**).

Therefore, these results suggest that despite having differences between baseline states, larval and adult progenitors respond to the spinal cord injury by adopting more similar gene regulatory programs with respect to their stage-specific context. Nevertheless, these injury-induced programs remain distinct from the embryonic progenitor state.

### Topic-based gene regulatory network inference distinguishes baseline and injury-induced gene regulatory programs in spinal cord progenitors

The observed changes in gene regulatory programs between iiERG and hERG cells were characterized using transcriptomic and epigenetic modalities separately. In addition, the activation of ERG cells might represent a continuous process, in which spinal cord progenitors respond to Tnf signaling (*27*), and acquire the injury-induced program to a varying degree. Therefore, to account for possible gradual shifts in GRNs and to determine whether the iiERG program is unique or shares regulatory features with other lesion-reactive cell states, we inferred GRNs with scDoRI (*15*) on a subset of the dataset containing injury-induced cell types (iiERG, iiOD and LRN) and their homeostatic counterparts (hERG, LFP, hOD and Neurons). scDoRI represents gene regulatory programs as latent topics, where each cell is modeled as a mixture of co-active topics rather than being assigned to a single discrete program. In our dataset, the inferred topics recapitulate the major cell states and separate both developmental stages and injury-induced identities (**Figure 2a,b**). ERG cell populations are primarily characterized by Topics 0, 19, 25, 39, and 41. While Topic 19 is shared across all developmental stages, other homeostatic topics exhibit stage-specific enrichment: Topic 0 is prominent in adult progenitors, whereas Topics 25 and 39 are associated with embryonic and larval progenitors. In contrast, Topic 41 is specifically linked to the injury-induced state. Low activity of Topic 41 in embryonic hERGs suggests that the injury response in larval and adult progenitors might involve a regulatory program that is not strongly presented during embryonic development. Therefore, we focused on the larval and adult-specific Topics 0, 19, and 41 (**Figure 2c**). Analogous homeostatic and lesion-associated topics were identified for oligodendrocytes (Topics 1 and 15) and neuronal lineages (Topics 29 and 48). In larvae, the injury-induced program of iiODs partially overlaps with that of iiERGs, suggesting both the close developmental proximity of oligodendrocytes and ERGs within the ventral pMN domain and spinal progenitors being oligodendrogenic but perhaps still able to be redirected towards regenerative neurogenesis after injury (*5*, *41*). In adults, the corresponding overlap is weaker, and Topic 1 shows higher activity in hODs than in larval ODs, probably due to a more mature oligodendroglial baseline and greater transcriptional separation from ERGs at adult stages. Therefore, the graded combinations of homeostatic and lesion-associated gene regulatory topics within cell identities suggests that spinal cord injury induces progressive regulatory remodeling rather than a binary switch between discrete cell states.

**Figure 2.**
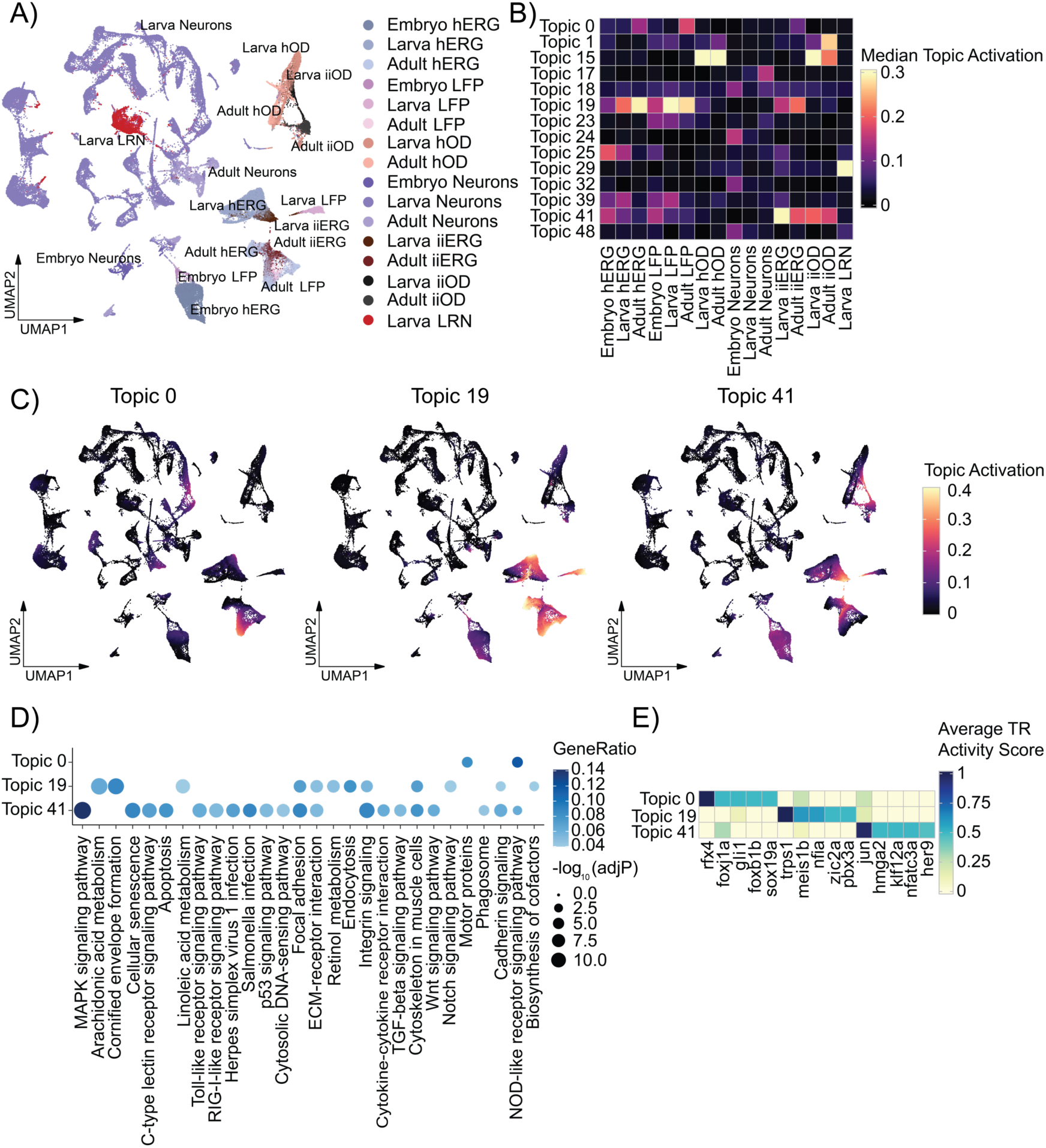
Topic-based GRN inference identifies homeostatic and injury-associated regulatory programs in spinal cord progenitors. (**A**) UMAP representation of the scDORI latent space for homeostatic and injury-associated cell types. (**B**) Median topic activity identifies cell type and developmental state-associated regulatory topics. Topics with median activity greater than 0.1 are shown. (**C**) Activity of homeostatic and injury-associated topics in cells selected for the GRN inference. (**D**) Results of the KEGG over-representation analysis of the top 300 genes associated with selected progenitor-related topics. (**E**) Average activity of top five upstream TRs associated with progenitor-related topics. hERG - homeostatic ependymo-radial glia; hOD - homeostatic oligodendrocytes; iiERG - injury-induced ERG; iiOD - injury-induced OD; LFP - lateral floor plate; LRN - lesion-reactive neurons.

To functionally characterize the gene regulatory programs represented by homeostatic and injury-associated topics in progenitor cells, we performed KEGG enrichment analysis on the 300 most strongly associated genes (**Figure 2d**). Homeostatic state-associated Topics 0 and 19 are enriched mainly for metabolic, ECM-related and a few signaling pathways such as Notch. The relatively low number of recovered KEGG pathways for Topic 0 could reflect a specific state of adult progenitors, which is reflected in the topic activity only in a subset of cells (**Figure 2c**). The iiERG-related Topic 41 shows enrichment for a variety of signaling pathways, with the strongest enrichment in the MAPK-signaling pathway. This result suggests the responsiveness of iiERG gene regulatory programs to exogenous cues after spinal cord injury (**Figure 2d**).

We next examined which inferred TRs constitute the ERG-related topics. Topics 0 and 19 are associated with distinct sets of TRs, indicating the developmental stage dependent divergence in the progenitor cells’ baseline state. Shared Topic 19 represents the MEIS/PBX developmental patterning program, while *nfia* implicates the presence of the gliogenic program in larvae and the quiescent-like program in adults (**Figure 2e**)(*34*, *36*). In turn, *gli1* activity in Topic 0 together with *zic* gene activity in Topic 19 suggests that the Hedgehog-associated program contributes to both larval and adult progenitor baseline states (*42–45*). Finally, the combination of *rfx4* and *foxj1a* indicates a distinct ciliated state of larval and adult ERG cells (*46*, *47*).

In contrast to the baseline-associated TRs of Topics 0 and 19, in the iiERG-associated Topic 41, the highest-ranking regulators include *jun*, *nfatc3a*, and *hmga2*. *hmga2* is known as a chromatin-associated regulator rather than as a classical DNA-binding TF. However, due to its ability to bind DNA and regulate gene expression, especially in neuronal progenitors, it was considered a TR (*48–51*). In addition, the presence of AP-1 subunit *jun* among the top regulators supports a stress-related and regeneration-associated identity of Topic 41(*27*). Therefore, hERGs at larval and adult stages appear to maintain distinct baseline regulatory states, whereas iiERGs converge on a rather specific injury-associated program.

### Topic-derived injury scores reveal continuous transitions of gene regulation toward lesion-reactive states

To quantify the observed continuum, we examined the distribution of progenitor cells along the ERG-associated topic axes across the time points (**Figure 3a**). Homeostatic ERG-associated Topics 0 and 19 broadly cover embryonic, larval and adult progenitors, whereas the injury-associated Topic 41 shows strongest activation in iiERGs. In larvae, Topic 41 activity peaks at 1-2 days post-lesion (dpl), concurrent with the active phase of regenerative neurogenesis(*5*). In adults, the separation between hERGs and iiERGs in injured animals along Topic 41 is less pronounced, suggesting that injury-induced regulatory remodeling is more gradual in adult progenitors. This difference in Topic 41 activation may reflect a stage-dependent baseline state of progenitor cells, with larval ERGs responding within a still developing spinal cord context(*5*), whereas adult cells are in a more quiescent state(*7*).

**Figure 3.**
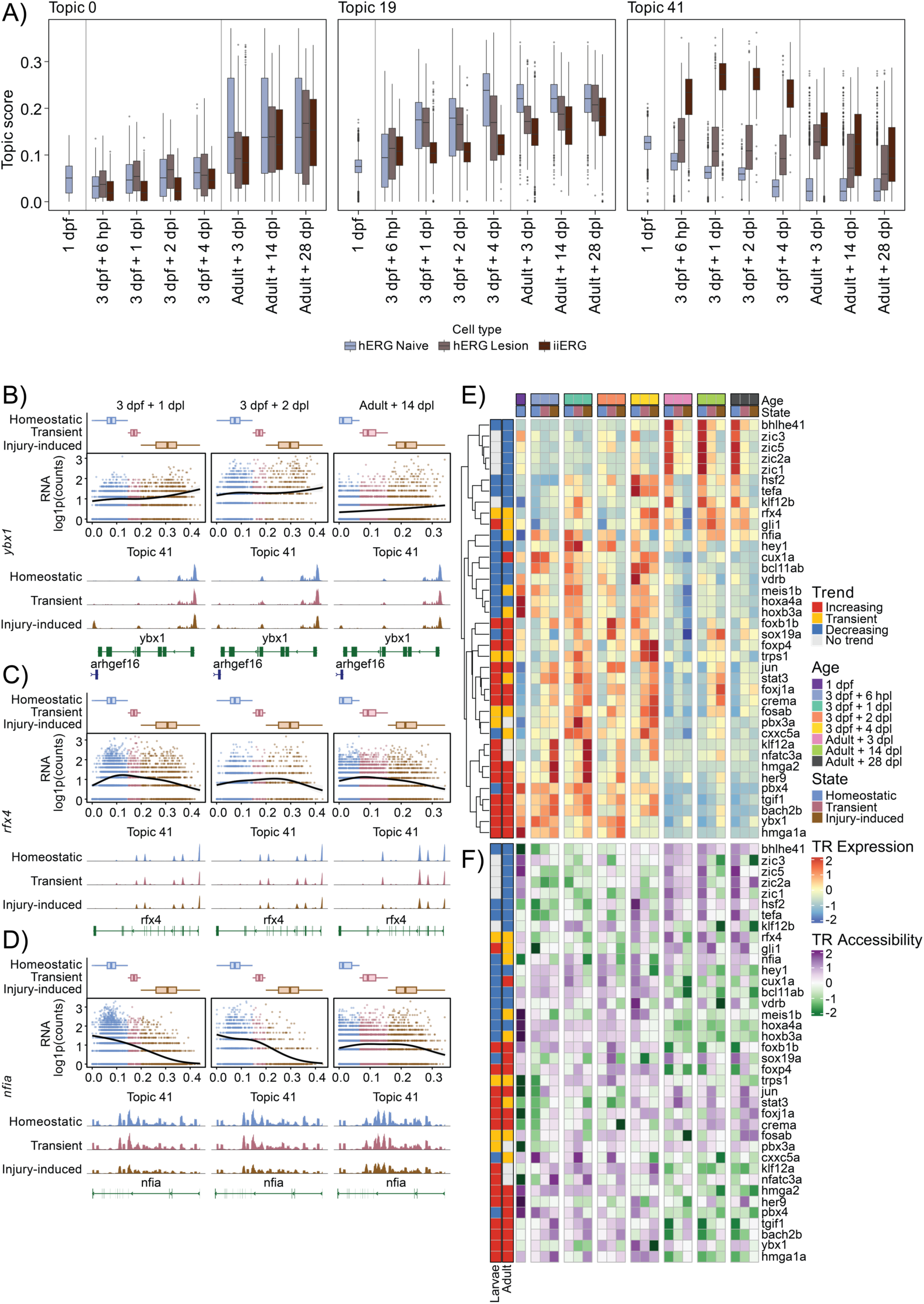
Injury-associated Topic 41 activity reveals a gradual shift from homeostatic to lesion-reactive ERG states. (**A**) Distribution of ERG cell states across the progenitor-associated Topics. Grey solid lines separate box plots by developmental stage. (**B**) Expression and accessibility profiles of *ybx1*. Top panel: Box plots showing distribution of cells along the Topic 41 axis across time points. Middle panel: *ybx1* expression increases along the injury-associated axis across developmental stages; bottom panel: chromatin accessibility profile of the gene body for *ybx1.* (**C**) Expression and accessibility profiles of *rfx4*. Top panel: Box plots showing distribution of cells along the Topic 41 axis across time points. Middle panel: Transient trend of *rfx4* expression along the injury-associated axis across developmental stages; bottom panel: chromatin accessibility profile of the gene body for *rfx4.* (**D**) Expression and accessibility profiles of *nfia*. Top panel: *nfia* expression decreases along the injury-associated axis across developmental stages; bottom panel: chromatin accessibility profile of the gene body for *nfia*. (**E**) Hierarchical clustering of Topic 0, Topic 19, and Topic 41 TRs average scaled expression across time points and cell states. (**F**) Aggregated promoter and gene body accessibility of Topic 0, Topic 19, and Topic 41 TRs across time points and cell states. Trend annotations and row order were transferred from expression data (Figure 3e). Genomic region 2 kb upstream of TSS was considered as a promoter. hERG - homeostatic ependymo-radial glia; iiERG - injury-induced ERG. dpf – days post fertilization, dpl – days post lesion, hpl – hours post lesion.

To investigate transcriptional changes along the injury-associated axis, we modeled raw gene counts as a function of Topic 41 using a negative binomial generalized additive model (NB-GAM). Using the first derivative of the fitted NB-GAM smooth with respect to Topic 41 score, we classified the modeled TRs into four expression trend classes: increasing, transient, decreasing and no trend (**Figure 3 b,c,d,e**). Since Topic 41 describes a gradient of regulatory programs across ERG cell states, we used logistic regression on Topic 41 scores to estimate the probability of an injury-induced state within each developmental stage. Based on this probability, cells were classified as Homeostatic (< 0.2), Transient (0.2–0.8), or Injury-induced (> 0.8) (**Supplementary Figure 13**).

Among the putative cell state regulators that increase with Topic 41, the most consistent TRs across stages are *foxj1a*, *jun*, *ybx1*, *hmga1a* and *hmga2*, while *stat3* shows a larval-specific upregulation trend after lesion, which is reflected at both transcriptomic and epigenetic layers (**Figure 3b, e, f**, **Supplementary Figure 15**). The upregulated module contains TRs that are shared with embryonic hERGs as well as TRs that are more restricted to iiERG states. *hmga1a*, *hmga2* and *ybx1* are highly expressed in embryonic hERGs and increase their expression with Topic 41 activity as larval and adult progenitors transition towards the injury-induced state, suggesting partial re-engagement of an embryonic progenitor-associated program in iiERGs. In contrast, *jun*, *stat3*, *foxj1a*, and *crema* are expressed more specifically in iiERGs indicating regulatory features specific to the lesion-reactive state. For example, *foxj1a*, a marker of ERG cells in zebrafish that is activated during spinal cord regeneration, shows a consistent increase in expression and gene-body accessibility (**Figure 3e, f**). An additional increase in expression of the Shh mediator *gli1* might also positively contribute to the *foxj1a* upregulation upon injury, since Shh signaling can activate *foxj1a* expression (*46*, *47*). In turn, *jun* upregulation reflects an AP-1-associated injury module activation, which is part of TNF/AP-1 signaling in zebrafish spinal progenitors and promotes regenerative neurogenesis (**Figure 3e, f**)(*27*).

The increase of chromatin-associated regulators *hmga1a*, *hmga2* and *ybx1* indicates the presence of a chromatin remodeling module during iiERG emergence. In particular, *hmga2* is a well-established regulator of neural stem-cell self-renewal, which suggests that the injury response partially re-engages a more plastic progenitor-like state (*50*). Despite low levels of expression in the adult injury-induced state, HMGA genes show an increase in chromatin accessibility at their gene bodies along the Topic 41 axis, which might indicate their activation (**Figure 3f**). In turn, the ability of *ybx1 to* mediate the activity of PRC2, which was shown in mouse embryonic stem cells, might additionally affect chromatin re-organization upon injury (*52*). The developmental TF *sox19a* also increases its expression in adult cells along the injury-induced axis. Its paralog *sox19b* was linked to EZH2-dependent histone methylation during zebrafish development(*53*), which might indicate that sox19a has a similar role in iiERGs. Therefore, the upregulated TRs define a regeneration-associated module that is shared across larval and adult stages. This module is acquired on top of distinct baseline states, accompanying a shift away from a gliogenic program in larvae and reactivation from a quiescent state in adults.

In larvae, injury-induced suppression of homeostatic regulatory programs is represented by downregulation of TRs such as *nfia* and *hey1* (**Figure 3d, e, f**, **Supplementary Figure 15**). *nfia* is one of the main regulators of the onset of gliogenesis in the developing spinal cord (*34*, *54*). The decline of a major Notch effector *hey1* suggests weakening of a homeostatic Notch-associated program in larval iiERGs (*55*). In adults, in addition to the downregulated Notch-associated module, *bhlhe41* and several *zic* genes also exhibit a downward trend in expression and accessibility, where *bhlhe41* is related to a quiescent-like program and *zic* genes, together with a transiently expressed *gli1,* to Shh signaling(*42–45*, *56*, *57*). Activation of the adult progenitor cells is further supported by the increase in cycling cells in the injury-induced state relative to the homeostatic state (**Supplementary Figure 16**). Thus, the declining module appears to represent the loss of a gliogenic maintenance program in larvae and loss of a quiescent homeostatic program in adults.

The transient regulators likely reflect the intermediate phase between baseline hERG identity and fully established iiERG state. *rfx4* shows a transient pattern in gene expression for both larvae and adults, while its gene body accessibility is decreased at the injury-induced state, showing a decline in accessibility prior to gene expression change (**Figure 3e,f**). The established role of RFX factors together with Foxj1 in ciliogenesis suggests temporary remodeling of the ciliated ependymal state during activation upon injury (*46*). In addition, the transient expression trend of *trps1* supports activation of a pro-regenerative program (*58*, *59*). However, our data do not allow us to determine whether the corresponding proteins remain active after transcript levels decline or whether protein activity decreases in parallel with transcription.

In summary, TR expression trends along the Topic 41-derived injury-associated axis reveal gradual remodeling of progenitor GRNs across developmental stages. This process involves both shared and stage-specific upstream regulatory factors, enabling larval progenitors to shift from a gliogenic baseline toward a regeneration-associated program, while adult ERGs appear to reactivate from a quiescent baseline state.

### Longitudinal gene set enrichment analysis refined the characterization of baseline-associated pathway changes in injury-induced progenitors

To characterize the temporal dynamics of lesion-reactive progenitor states, we performed longitudinal differential expression analysis comparing injury-induced and homeostatic ERGs across time points. DEGs were ranked by average log2 fold change and subjected to KEGG-based Gene Set Enrichment Analysis (GSEA) to identify time point-specific pathway changes.

In larval progenitors, the injury-associated program is characterized by sustained enrichment of extracellular signaling and cytoskeletal remodeling pathways across time points, including Integrin signaling, Focal adhesion, and MAPK- and TGF-beta-associated pathways (**Figure 4**). In parallel, the Notch signaling pathway shows negative enrichment in larval iiERGs relative to hERGs at 6 and 24 hpl. In contrast, adult iiERGs show weaker enrichment of cytoskeletal remodeling pathways and stronger representation of biosynthetic programs, including protein processing in the endoplasmic reticulum and aminoacyl-tRNA biosynthesis. Neuroactive and metabolic pathways show significant negative enrichment in the adult injury response.

**Figure 4.**
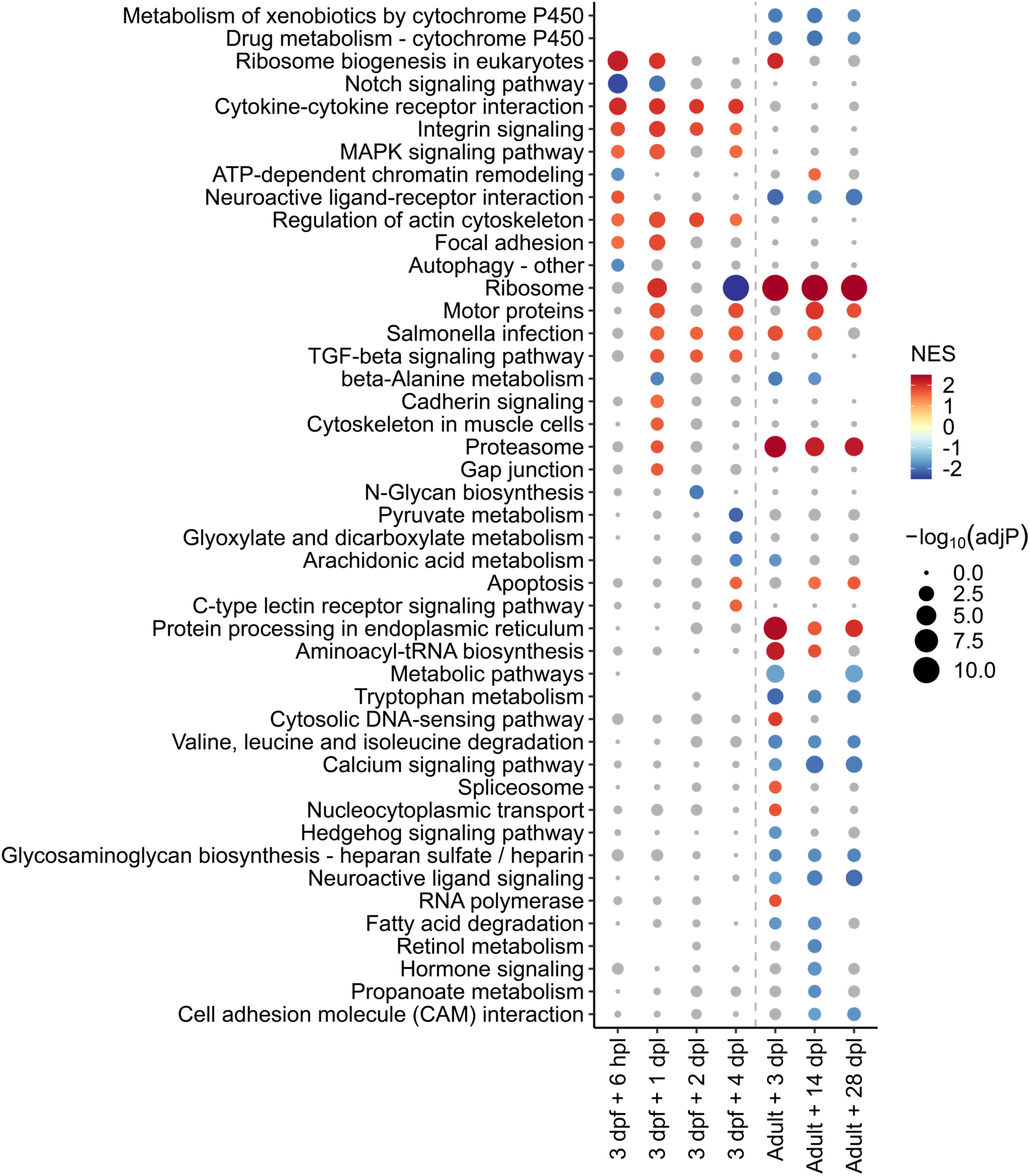
Longitudinal KEGG pathway GSEA of DEGs between homeostatic and injury-induced states of ERG cells reflects the temporal dynamics in the regulatory programs of the progenitor cells after injury. dpf – days post fertilization, dpl – days post lesion, hpl – hours post lesion.

The observed pathway-level differences refine the shared injury-associated regulatory program identified by topic and TR analyses. Since KEGG GSEA compares injury-induced and homeostatic ERGs within each developmental stage, enriched pathways are expected to reflect the injury response in the context of stage-specific baseline states. For example, the gliogenic Notch pathway is negatively enriched in larval iiERGs(*60*), while adult iiERGs show broader metabolic remodeling. Thus, larval and adult iiERGs show distinct pathway signatures despite sharing a common injury-associated regulatory core marked by Topic 41 activity, e.g., induction of *jun*, *hmga1a*, and *hmga2* (**Figure 3e**).

In summary, these findings support a model in which spinal cord injury induces stage-dependent regulatory programs. Larval progenitors enter a remodeling and neurogenesis-permissive state, whereas adult progenitors undergo a more gradual transition coupled to exit from a quiescent homeostatic program (**Figure 5**). This suggests that larval and adult iiERGs acquire a related lesion-reactive identity through different regulatory routes.

**Figure 5.**
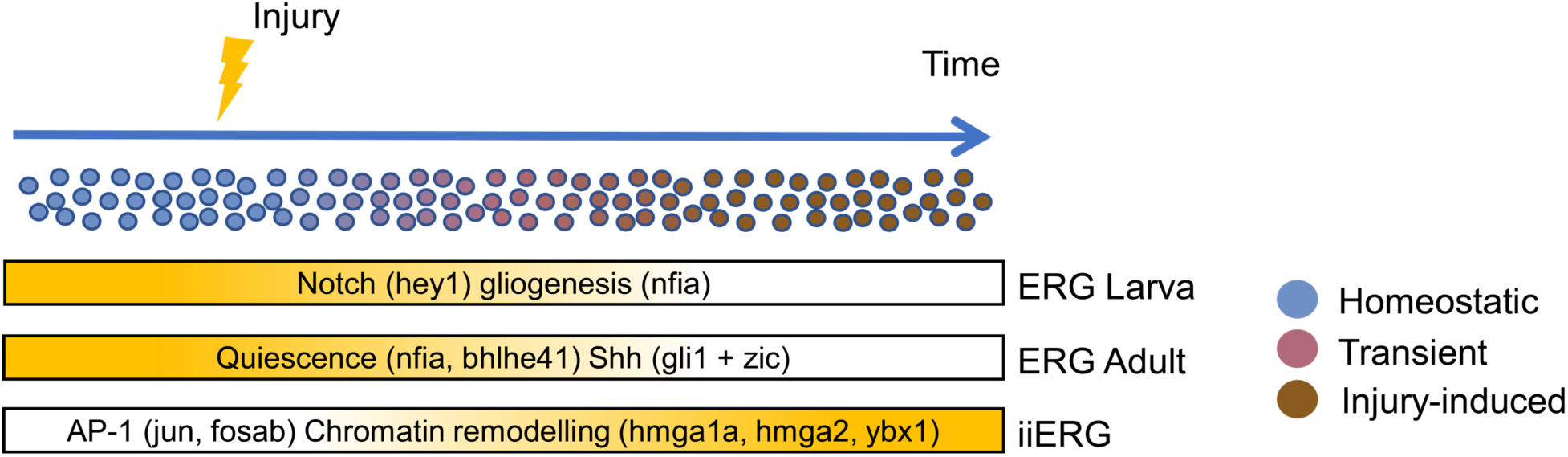
Developmental stage-specific baseline regulatory programs in ERG cells shift to the common injury-induced state after the spinal cord injury. ERG - ependymo-radial glia; iiERG - injury-induced ERG.

## Discussion

This study defines gene regulatory transitions in injury-induced activation of zebrafish spinal cord ERGs as a continuous regulatory transition and shows that larval and adult ERGs converge on a related injury-induced state through partially shared, but non-identical, regulatory routes. Larval and adult baseline cellular states in the spinal cord differ substantially: despite showing low neurogenic activity, larval tissues are still developing, whereas adult hERGs exist in a comparatively quiescent state. These differences likely influence how regenerative gene regulatory programs can be activated. Topic-based analysis showed that larval ERGs display a sharper separation in their gene regulatory programs along the injury-associated axis, whereas adult ERGs undergo a more gradual redistribution. Importantly, the continuous transition from the baseline to the injury-induced gene regulatory state is marked by the activation of a set of TRs that is independent of the developmental stage.

The divergence between homeostatic programs in larval and adult hERGs is more apparent in the declining regulatory modules than in the acquired injury-induced cell state, which is mostly shared between developmental stages. In larvae, these modules reflect a gliogenic and Notch-associated baseline, whereas in adults they align more closely with a quiescent homeostatic identity. *nfia,* which shows a downward expression trend at both stages, drives gliogenesis in larvae but likely supports the quiescent cell state in adults, as demonstrated for adult Müller glia in the zebrafish retina(*36*). Additional support for an exit of larval ERGs from gliogenesis is provided by the upregulation of *lin28a* (**Figure 1c**), which has been shown in a mouse model to block gliogenesis and promote neurogenesis (*61*). Transient expression of *trps1* might further indicate the decline of the Notch program in progenitor cells by inhibiting *sox9* and therefore the onset of the pro-regenerative program (*41*, *58*, *59*, *62*). In adult ERGs, reduced expression of Notch-linked *nfia* and *hey1* in combination with *zic* genes further suggests downregulation of the Notch pathway, which could lead to the activation of regenerative neurogenesis(*43*, *55*, *63*). In addition, Zic proteins have been shown to interact with Gli proteins, thereby modulating Shh signaling (*42*, *44*, *45*). This may indicate that a temporary downregulation of Shh is required for adult ERGs to exit the quiescent state, particularly because prolonged Shh signaling has been linked to the accumulation of quiescent murine neural stem cells (*64*). However, *shha* was shown to be upregulated in the zebrafish adult spinal cord after injury and inhibition of Shh signaling led to decrease in ventricular proliferation and regenerative neurogenesis(*65*). Together, these results suggest that Shh signaling is likely fine-tuned in a time-specific manner during the adult injury response.

The metabolic and translational remodeling in adult iiERGs might be also related to exit from quiescence. In other progenitor systems, inhibition of mitochondrial pyruvate import promotes activation of adult neural stem cells (*66*), while *Spot14* knockdown increases their proliferation by decreasing a suppression of Fasn-dependent lipogenesis (*67*). Another example is muscle satellite cells, whose quiescence exit is accompanied by increase in translation activity (*68*).

The presence of AP-1 subunits within Topic 41 suggests that adult progenitors respond to macrophage-derived TNF in a manner similar to larval ERGs(*27*). This activation might be linked to a broader process of chromatin and cell-state remodeling. Specifically, HMGA-family factors have been implicated in zebrafish Müller glia reprogramming, where *hmga1a* is induced in reactive glia and is required for the transition to a proliferative regenerative state(*36*). Consequently, *hmga1a* and *hmga2* may act through a similar mechanism in larval and adult spinal cord progenitors (*49*, *50*). Furthermore, the interaction between *ybx1* and PRC2 activity in iiERGs might create a permissive environment for neuronal commitment, mirroring mechanisms reported in mouse embryonic stem cells (*52*). In larvae, additional support for PRC2 fine-tuning might be provided by the upregulated expression of *sox19a*. *sox19a* and *sox19b* belong to a functionally redundant B1 Sox group (*69*), and *sox19b* was shown to promote neural progenitor proliferation and limit premature neuronal differentiation through EZH2-mediated histone methylation(*53*). Therefore, *sox19a* might exhibit a similar function in iiERG cells. Thus, the emergence of iiERGs requires both acquisition of an injury-associated module and erosion of the opposing baseline program.

While GRN inference provides insights into regulatory modules altered after injury, current approaches remain largely correlative and therefore require cautious interpretation. Recent work has argued for a shift toward mechanistically constrained representation-learning models that infer latent regulatory features rather than dense correlation-based networks (*70*). scDoRI partly addresses this by modeling cells as mixtures of enhancer-linked topics under explicit regulatory constraints, which allows cluster-free analysis of continuous regulatory variation (*15*). However, the identified TRs still remain candidate regulatory programs that require orthogonal evidence. In parallel, perturbation-based interpretation is complicated because many of them likely act in combination, in multiple cellular processes, and may be compensated for by other mechanisms. In addition, since our sample preparation was designed to enrich for spinal cord progenitors, we cannot exclude that intermediate or descendant states of lesion-reactive neurons and ODs may have been underrepresented or missed. Therefore, a full characterization of these lineages will require additional unsorted samples.

In conclusion, we propose that spinal cord regeneration in zebrafish engages ERGs through a staged and continuous process of remodeling gene regulatory programs. A shared lesion-associated module, focused on stress-responsive and chromatin-remodeling regulators, is activated in both larvae and adults, yet it is implemented on top of distinct, stage-specific baseline programs. Thus, iiERGs represent a regeneration-specific state achieved through the gradual regulatory remodeling of pre-existing hERG identity. Our atlas of regenerative neurogenesis in the zebrafish spinal cord across developmental stages reveals core gene-regulatory mechanisms that facilitate regenerative neurogenesis regardless of developmental stage that could be targeted in non-regenerating vertebrates in the future.

## Methods

### Animals

Tg (*her4.1:eGFP*)^y83Tg^ line, abbreviated as *her4.1:eGFP*(*16*), was raised and kept under standard conditions (*71*). Zebrafish experiments were performed under the state of Saxony license TVV 18/2024, and holding licenses DD24-5131/364/11, DD24-5131/364/12.

### Spinal cord lesion in larvae

Lesions of larvae were performed as previously described(*5*). Briefly, at 3 dpf, zebrafish larvae were deeply anaesthetized in E3 medium (*72*) containing 0.02 % of MS-222 (Sigma). Larvae were then transferred to an agarose plate. After removal of excess water, larvae were placed in lateral positions. A transection of the entire spinal cord was made using a 30 G syringe needle at the level of the 15th myotome, without injuring the notochord.

### Cell dissociation from larvae

Cell dissociation from embryos and larvae was performed as described previously(*17*). In brief, using the Tg(her4.1:eGFP) line, we excised the whole trunk from 1 dpf embryos (110 embryos), or the trunk surrounding the injury site from larvae, pooling 225 larvae per condition. GFP^+^ cells (125,000 per condition) were collected by fluorescence-activated cell sorting (FACS). The sorted cells were pelleted at 500 × g for 5 min at 4 °C, and the supernatant was removed to leave a final volume of approximately 50–100 µL. Nuclei were then extracted as described below.

### Cell dissociation from adult fish

Spinal cord injury was performed in approximately 30 adult fish (6-8 months old) as described previously(*18*). The spinal cord was dissected three vertebrae rostral and three vertebrae caudal to the injury site. Samples were collected in cold 1× PBS and kept on ice until all fish had been processed; in total, 20–22 adult fish were processed within approximately 2 h. The PBS was then removed, and the larval dissociation protocol described above was applied, with the addition of 30 µL of Myelin Removal Beads II (Miltenyi Biotec, Cat# 130-096-733). Samples were triturated with a flame-polished glass pipette at room temperature (∼22–23 °C). Following complete dissociation (∼5 min), the 2 mL suspension was split into two 2 mL Eppendorf tubes (1 mL each), and 1 mL of DFP buffer (DMEM, FBS, protease inhibitor) was added to each and mixed thoroughly. Cells were passed through the magnetic (MACS) column within approximately 1 min and then centrifuged immediately. Viable GFP⁺/PI⁻ cells were isolated by FACS; for adult fish, approximately 190,000–210,000 GFP⁺ cells, in total, were recovered. The sorted cells were pelleted at 500 × g for 5 min at 4 °C, and the supernatant was removed to leave a final volume of approximately 50–100 µL. Nuclei were then extracted as described below.

### Nuclei extraction for 10x Multiome (ATAC and Gene Expression)

The supernatant was carefully removed, leaving approximately 50–100 µL, and 1 mL of nuclei extraction buffer containing 0.2 U/µL RNase inhibitor was added. For adult fish, we additionally added 10µL of 10% NP-40 to the buffer. The suspension was pipetted gently 10 times over 1 min, followed by a 1 min incubation on ice; this cycle was repeated five times and was followed by centrifugation at 500 × g for 5 min at 4 °C.

The supernatant was removed by pipetting, and the pellet was washed with 1 mL of wash buffer (840 µL nuclease-free H₂O, 10 µL Tween-20, 10 µL 1 M Tris-HCl pH 7.4, 2 µL 5 M NaCl, 3 µL 1 M MgCl₂, and 135 µL 7.5% BSA), followed by centrifugation at 500 × g for 5 min at 4 °C. The supernatant was removed, and the pellet was resuspended in 1× DNB buffer (7.5 µL 1× DNB buffer, 142.6 µL nuclease-free H₂O, 1.5 µL 40 U/µL RNase inhibitor) and centrifuged again. The supernatant was removed to leave approximately 3 µL, and the pellet was resuspended and brought to a final volume of 6 µL. Nuclei quality was assessed visually under a 20× objective, and the concentration was determined from a 1:10 dilution in 1× PBS using a Neubauer counting chamber. Approximately 16,000 nuclei were used as input for the 10x Multiome kit.

### Single-nuclei multiome sequencing: ATAC + Gene Expression

Open chromatin within the nuclei was tagmented for 1 h at 37 °C using the Chromium Next GEM Single Cell Multiome ATAC Kit A (PN-1000280), according to the Chromium Next GEM Single Cell Multiome ATAC + Gene Expression protocol (Document no. CG000338, Rev F/G, 10x Genomics). During this step, adaptor sequences were appended to the ends of the DNA fragments.

The tagmented reaction was then processed to generate cell-barcoded, full-length cDNA and ATAC libraries. Reverse transcription and barcoding reagents from the Chromium Next GEM Single Cell Multiome Reagent Kit A (PN-1000282) were added directly to the tagmented reaction. This mixture was loaded onto a Chromium Next GEM Chip J and processed according to the Chromium Next GEM Single Cell Multiome ATAC + Gene Expression protocol (Document no. CG000338, Rev F/G, 10x Genomics). GEMs were generated to produce 10x-barcoded transposed DNA and 10x-barcoded full-length cDNA. After quenching the reaction with the reagent provided by 10x Genomics, the emulsion was broken and the barcoded products were purified using silane magnetic beads.

To obtain sufficient material for library construction and to fill gaps, the barcoded products underwent a 7-cycle pre-amplification, after which they were purified with 1.6× SPRI beads and eluted in 160 µL of Elution Buffer (Qiagen). Of the pre-amplified product, 40 µL was used to construct sequencing-ready ATAC libraries via an 8-cycle index PCR, and 35 µL was used to specifically amplify cDNA in an 8-cycle PCR. The ATAC libraries underwent double-sided size selection with SPRI beads, and the amplified cDNA underwent a 0.6× SPRI purification.

Gene Expression libraries were generated from 25% of the cDNA material using the 10x Genomics Library Construction Kit B (PN-1000190), comprising fragmentation, dA-tailing, adapter ligation, and a 12-cycle indexing PCR, according to the manufacturer’s protocol. Both the ATAC and Gene Expression (GEX) libraries were assessed for quality and quantity. The libraries were sequenced on an Illumina NovaSeq 6000 in paired-end mode (R1/R2: 100 cycles; I1/I2: 10 cycles), generating more than 70 million fragment pairs.

### Genome alignment

We used the GRCz11 assembly of the zebrafish genome (Ensembl release 109). GFP, dsRed, and mCherry sequences were added to both the FASTA and GTF files. The modified FASTA and GTF files were used as input to the ‘cellranger-arc (v2.0.1) mkref’ command to build a custom reference. The FASTQ files were then aligned with ‘cellranger-arc count’.

### Quality control of snRNA-seq and snATAC-seq data

Single-nuclei multi-omic data were processed using Seurat (v5.0.3) (*73*) for snRNA-seq and Signac (v1.13.0) (*74*) for snATAC–seq. For each sample, the percentage of the RNA-seq reads mapped to the mitochondrial genes and nucleosome signal, Transcription Start Site (TSS) enrichment and Fraction of Reads in Peaks (FRiP) were calculated. Cells that passed the following threshold were retained: RNA unique molecular identifier (UMI) between 500 and 8000, ≥300 detected genes, mitochondrial RNA percentage ≤15%, ATAC counts between 1000 and 30000, TSS enrichment ≥4, and nucleosome signal ≤2. To prevent exclusion of otherwise high-quality but lower-depth ATAC cells, an ATAC “rescue” strategy was applied when ATAC counts were below the main cutoff: cells with ATAC counts <1000 were kept only if they had TSS enrichment ≥4, FRiP ≥0.8, and ATAC counts ≥500. The doublets were estimated using non-filtered data. The scDblFinder (v1.16.0) (*75*) was separately applied for the RNA and ATAC modalities. For snATAC-seq the combination of two doublet calling approaches was used: the original scDblFinder methods (aggregateFeatures = TRUE, nfeatures = 50, processing = “normFeatures”) and the scDblFinder adaptation of the Amulet method. Afterwards, the p-values from both doublet calling methods were aggregated using the sumlog function of the metap library. Cells that were identified as doublets in both modalities were removed.

### snRNA-seq normalization and feature selection

RNA normalization was performed per sample using the pooling-based size factor estimation strategy. First, mitochondrial genes were removed from the count matrices, and genes which were expressed in less than 20 cells within each sample were filtered out. Next, cells were pre-clustered with the quickCluster function from the scran library (v1.30.2) (*76*), and those groups were used for the pooling-based size factor estimation. Afterwards, log-normalized with the pseudocount of 1 expression values were computed. Per-sample log-normalized matrices were then concatenated across samples. Raw RNA counts were used for Highly Deviant Genes (HDGs) identification. Deviance-based feature selection using scry library (v1.14.0) (*77*) was applied for each sample. Per-sample deviance rankings were aggregated by median rank across samples and the top 3000 genes were selected as variable features. These features were centered and scaled, and Principal Component Analysis (PCA) was run on the scaled HDGs.

### snATAC-seq normalization and feature selection

ATAC peaks were annotated using the Ensembl release 109 via the AnnotationHub library. For the downstream analysis all the alternative loci as well as the mitochondrial DNA were removed. Per-sample peak sets were merged into a consensus peaks set and ATAC counts were quantified for all cells against the resulting peaks set. The feature matrix was normalized using Term Frequency-Inverse Document Frequency (TF-IDF) normalization. The variable features were selected with the FindTopFeatures function from the Signac library (*74*). The resulting set of features was used for the Singular Value Decomposition (SVD) on the TF-IDF-transformed matrix.

### Batch correction and integration of RNA and ATAC modalities

To account for the batch effect across samples, Harmony (v1.2.0) (*78*) was independently applied to the top 30 principal components for snRNA-seq data and top 30 LSI dimensions for snATAC-seq (excluding the first dimension). The corrected embeddings were integrated using Weighted Nearest-Neighbor (WNN) analysis(*79*). Clustering was performed on the WNN graph using the SLM clustering algorithm(*80*) with a resolution of 0.3. Uniform Manifold Approximation and Projection (UMAP) (*81*) was generated for WNN embeddings using the RunUMAP function with default parameters from the Seurat library (*73*).

### Cell type annotation

Cell-type annotation was performed using a list of marker genes (Supplementary Table 1). To account for developmental stage-specific differences, cells were split into three groups: embryonic (1 dpf), larval (3-7 dpf) and adult samples. The same pre-processing approach was applied for each stage as was used for the complete dataset: RNA HDGs were re-selected using the same deviance/median-rank approach, ATAC LSI was recomputed, batch correction with Harmony (*78*), WNN neighbors(*79*), clustering and UMAP embeddings were rerun stage-wise. Subclustering was performed by subsetting selected SLM clusters and reintegrating the subset as described above.

### Cell type-specific peak calling

Cell type-specific peaks were called with MACS v3.0.3(*82*) using the CallPeaks function from the Signac library (*74*). The newly generated set of peaks was processed as it was described in the snATAC-seq normalization and feature selection section.

### Identification of cell type-specific motif activity and expression of corresponding TFs

A zebrafish-specific motif database was retrieved from the Catalog of Inferred Sequence Binding Preferences (CIS-BP) database version 3 (*83*). To reduce the motif redundancy, we followed the strategy used in the MEME database for the construction of non-redundant motifs set for the CIS-BP database version 2. If the direct motif was present, it was initially selected, otherwise the inferred motif with the highest DNA binding domain similarity according to CIS-BP was used. When there was more than one direct or inferred motif for a given TF with the same DNA binding domain similarity, the selection was made according to CIS-BP’s “Motif_Type” attribute. The motif discovery methods were ordered in the following manner: ChIP-seq+ChIP-exo, ChIP-exo, ChIP-seq, Dap-seq, HocoMoco, DeBoer11, PBM, SELEX, SMiLE-seq, B1H, High-throughput SELEX SAGE, PBM:CSA:DIP-chip, ChIP-chip, EMSA, COMPILED, DNaseI footprinting. Motifs that were labeled Transfac, Misc or Unknown were not considered. Resulting Position Frequency Matrices (PFMs) were converted into Position Weight Matrices (PWMs) using the universalmotif library (v1.26.2) (*84*) and used for downstream analysis. The set of non-redundant PWMs was used to infer the motif activity score utilizing chromVAR (*85*) with the default parameters.

### Identification of cell type-specific motif activity and TF expression

The wilcoxauc function from the presto (v1.0.0) library (*86*) was used to obtain the cell type-specific motif activity scores and TF expression values. The average of resulting Area Under the Receiver Operator Curves (AUCs) for RNA and ATAC modalities was used to select the cell type-specific TF motif activities.

### Gene Regulatory Network analysis

Raw paired gene expression and chromatin accessibility matrices from injury-induced and homeostatic progenitor cells, oligodendrocytes and neurons were used as input for scDoRI (*15*). The scDoRI workflow was adapted in two respects: first, ATAC quantification during metacell construction and peak selection was modified to handle fragment-based counts rather than applying the default insertion-to-fragment conversion; second, L1/L2 regularization was implemented directly on trainable model parameters during optimization. Protein-coding genes were retained based on Ensembl release 109 and TFs were defined by the presence of a corresponding CIS-BP motif. Before feature selection, genes expressed in at least 5% of cells in any cell type at any time point were retained; from these, 5,500 highly variable genes and up to 500 highly variable TFs were selected, yielding 468 TFs in the final GRN analysis. Metacells were constructed from RNA-based clustering after normalization, log-transformation, PCA and Harmony correction. A final set of 150,000 ATAC peaks was retained by first keeping all promoter peaks and then selecting the remaining non-promoter peaks ranked by the standard deviation of mean accessibility across metacells. Candidate *cis*-regulatory peaks were defined within ±75 kb of each gene, and peak-by-motif scores were computed from 500-bp peak-centered sequences using FIMO (*87*). scDoRI was run with 50 topics, a maximum of 70 epochs for the GRN inference step and the patience parameter for the early stopping of 10 epochs for all training steps, while all remaining parameters were kept at default settings. Topic regulators were identified using the composite TF activity score (*15*). In brief, TF-to-gene weights from the final topic-specific GRNs were summed across target genes and normalized by the total weight within each topic. The composite TF activity score was then calculated as the mean of TF intensity and TF specificity. TF intensity represents the relative activity of a TF compared with other TFs within the same topic, whereas TF specificity represents the relative activity of the same TF in one topic compared with its activity across all topics. TFs with a score > 0.05 were defined as topic regulators.

### KEGG over-representation analysis of topic-associated genes

For selected topics, the top 300 topic-associated genes with the highest score were used for KEGG (*88*) over-representation analysis. Pathway enrichment was performed with the enrichKEGG function of the clusterProfiler library (*89*) using the zebrafish annotation (organism = “dre”). P-values were adjusted using a Benjamini–Hochberg multiple-testing correction. Pathways supported by fewer than five genes were excluded.

### Negative Binomial Generalized Additive Model fitting

Gene expression changes along the selected topic were modeled using a negative binomial generalized additive model implemented in the mgcv library (*90*). Raw RNA counts were used as input, and the corresponding topic’s score was included as a continuous predictor using a cubic regression spline. The differences in sequencing depth were controlled by an offset based on centered log library size factors, and each sample was modeled as a random effect. Models were fitted with the bam function from the mgcv library (*90*) using a log link. To minimize a confounding effect between injury response and larval maturation, comparisons were made relative to age-matched non-lesioned controls whenever available. Thus, for this and subsequent analyses, in larvae, the 3 dpf Naive and 3 dpf + 6 hpl Lesion samples were combined, as a Naive control was absent for the 6 hpl time point and developmental change was considered negligible over 6 hours. For adult fish, the Adult samples were grouped with all Lesion time points and considered to be a stable homeostatic baseline.

### Assignment of expression trends from fitted NB-GAMs

Expression trends for modeled TRs along an injury-associated topic were assigned using the first derivative of each fitted NB-GAM smooth with respect to the topic score. Derivatives were estimated with the gratia package (*91*) on a grid of 200 points using central finite differences and simultaneous confidence intervals. At each grid point, the derivative was classified as increasing when the lower confidence bound was greater than zero, decreasing when the upper confidence bound was less than zero, and flat otherwise. To account for local fluctuations, directional segments shorter than or equal to 20 consecutive grid points were reclassified as flat. After removal of flat segments, the remaining sign pattern was used to assign one of four trend classes to each fitted model: increasing, transient, decreasing or no trend. Increasing and decreasing trends corresponded to a single sustained positive or negative derivative segment, respectively, whereas a transient pattern corresponded to up-then-down or down-then-up transitions. Models without a sustained non-flat segment were classified as no trend. To summarize trends across time points, trend assignments were aggregated within larval and adult developmental stages. A consensus trend for each TR was defined as the most frequently observed trend category across modeled age panels within a developmental stage. Ties were resolved using the following priority order: increasing, decreasing, transient and no trend.

### Definition of cell states using logistic regression on Topic scores

Cell states were defined for selected cell types using a binomial logistic regression model. For each developmental stage, topic scores from the corresponding injury-induced cell population were used to train the model to distinguish homeostatic and injury-induced cell states. Prior to model fitting, topic scores were centered and scaled within each developmental stage using the corresponding training subset. The trained model was then applied to all cells of the respective cell type to estimate the probability of injury-induced cell state. Cells with predicted probability < 0.2 were classified as Homeostatic, cells with predicted probability > 0.8 were classified as Injury-induced, and cells with intermediate probabilities were assigned to the Transient state.

### Cell-cycle score annotation across Topic-defined ERG states

S and G2/M human cell cycle genes (*92*) were mapped to zebrafish orthologs using the biomaRt (v2.58.0) library (*93*, *94*). Cell-cycle module scores were calculated for S and G2/M cell cycle phases using gene sets mapped by orthology with the AddModuleScore function from the Seurat library (*73*). Cells were assigned to either S or G2/M phase according to the higher of the two phase-specific module scores, while cells with scores less than or equal to zero were classified as Non-cycling. Cell-cycle proportions were summarized at the sample, time point and cell state level. Groups with fewer than 50 cells were excluded from the analysis. The fraction of cycling cells was defined as the number of S-phase and G2/M cells divided by the total number of cells in the corresponding group. For each time point, a linear model was fitted with the sample-state-level fraction of cycling cells as the response variable and the median Topic 41 score as the predictor. Spearman correlation coefficient between median Topic 41 score and the fraction of cycling cells was additionally calculated. P-values from time point-specific Spearman rank-correlation tests were adjusted using the Benjamini–Hochberg method.

### Differential analysis

Differential gene expression, accessibility, and motif activity analyses were performed using a sample-level aggregation strategy. Profiles containing fewer than 50 cells were excluded. Differential gene expression and accessibility analyses were performed using the DESeq function from the DESeq2 library (v.1.46.0) (*95*) with default parameters. Differential motif activity analyses were performed using the lmFit, contrasts.fit, and eBayes functions from the limma library (v.3.62.2) (*96*). When only one replicate was available in each group, the comparison was performed using the FindMarkers function from the Seurat library (*73*).

For differential gene expression analysis, raw RNA counts were summed across cells within each pseudobulk sample. Genes with fewer than 10 total counts across all pseudobulk samples included in a comparison were excluded.

For gene-level differential accessibility analysis, ATAC counts from peaks were summarized over gene bodies and their 2 kb upstream promoter regions in a strand-aware manner. Gene coordinates were obtained from Ensembl release 109 through the AnnotationHub library. A peak-to-gene mapping matrix was used to aggregate peak counts into gene-level accessibility counts for each pseudobulk sample. Genes with fewer than 10 total accessibility counts across all pseudobulk samples included in a comparison were excluded.

For differential chromVAR analysis, chromVAR deviation z-scores were averaged across cells belonging to the same biological replicate and comparison group.

For analyses performed using DESeq2 or limma, p-values were adjusted for multiple testing using the Benjamini–Hochberg procedure. For comparisons performed using the FindMarkers function, p-values were adjusted using the Bonferroni correction. DEGs, DARs, and differentially active motifs were considered significant at the adjusted p-value < 0.05. For differential expression and accessibility analyses, features were additionally required to have an absolute log2 fold change > 0.1. Features with positive log2 fold change were required to be detected in at least 5% of cells in the tested group, whereas features with negative log2 fold change were required to be detected in at least 5% of cells in the reference group.

### KEGG pathway enrichment analysis

KEGG (*88*) pathway enrichment was performed using the differential expression results. Gene symbols were mapped to Entrez identifiers using the org.Dr.eg.db database (v3.18.0), and genes that could not be mapped were excluded from the KEGG analysis.

Preranked KEGG gene-set enrichment analysis (GSEA) (*97*, *98*) was performed using the gseKEGG function from the clusterProfiler library (v4.8.3) (*89*). Genes were ranked by log2 fold change. Genes with positive log2 fold change were required to be detected in at least 5% of injury-induced cells, whereas genes with negative log2 fold change were required to be detected in at least 5% of homeostatic cells. When multiple gene symbols mapped to the same Entrez identifier, the entry with the largest absolute log2 fold change was retained. P-values were adjusted using the Benjamini–Hochberg multiple test correction and pathways with adjusted p-value < 0.05 were considered significant.

## Data availability

Data generated in this study (10x Multiome (ATAC and Gene Expression)) have been deposited in GEO and will be made available upon peer review/publication.

## Code availability

The full code is available at Zenodo (https://doi.org/10.5281/zenodo.21474412).

## Acknowledgments

This work was supported by the Center for Scalable data analytics and artificial intelligence (Scads.AI) Dresden-Leipzig. The experiments, **DS**, and **MIC** were funded by the German Research Foundation (DFG) project 514187677. **ARP** was supported by the Mildred Scheel Early Career Center Dresden P2, funded by the German Cancer Aid. **CGB** was supported by an Alexander von Humboldt Stiftung Professorship award and TU Dresden core funding. This work was supported by the Light Microscopy Facility, the DRESDEN-concept Genome Center, the Flow Cytometry Facility, the Zebrafish Facility and all core facilities of the Center for Molecular and Cellular Bioengineering (CMCB) at the Technische Universität (TU) Dresden. We are grateful to the whole Biomedical Genomics group at TU Dresden for fruitful discussions and feedback, and to Fabian Rost, Daniel Wehner, and Andreas Beyer for valuable input on the manuscript. We additionally thank Michaela Unger for providing illustration of the experimental design.

## Contributions

**ARP, CGB,** and **TB** conceptualized the study. **JR** helped in the conceptualisation of the experimental design. **MIC**, **SR**, **MW**, **KH**, **DB**, and **AB** collected and prepared the data. **DS** and **MIC** performed data analysis, supervised by **ARP**. **DS** and **ARP** wrote the manuscript with help from **MIC** and input from all authors.

## Supplementary Material

**Supplementary Table 1.**
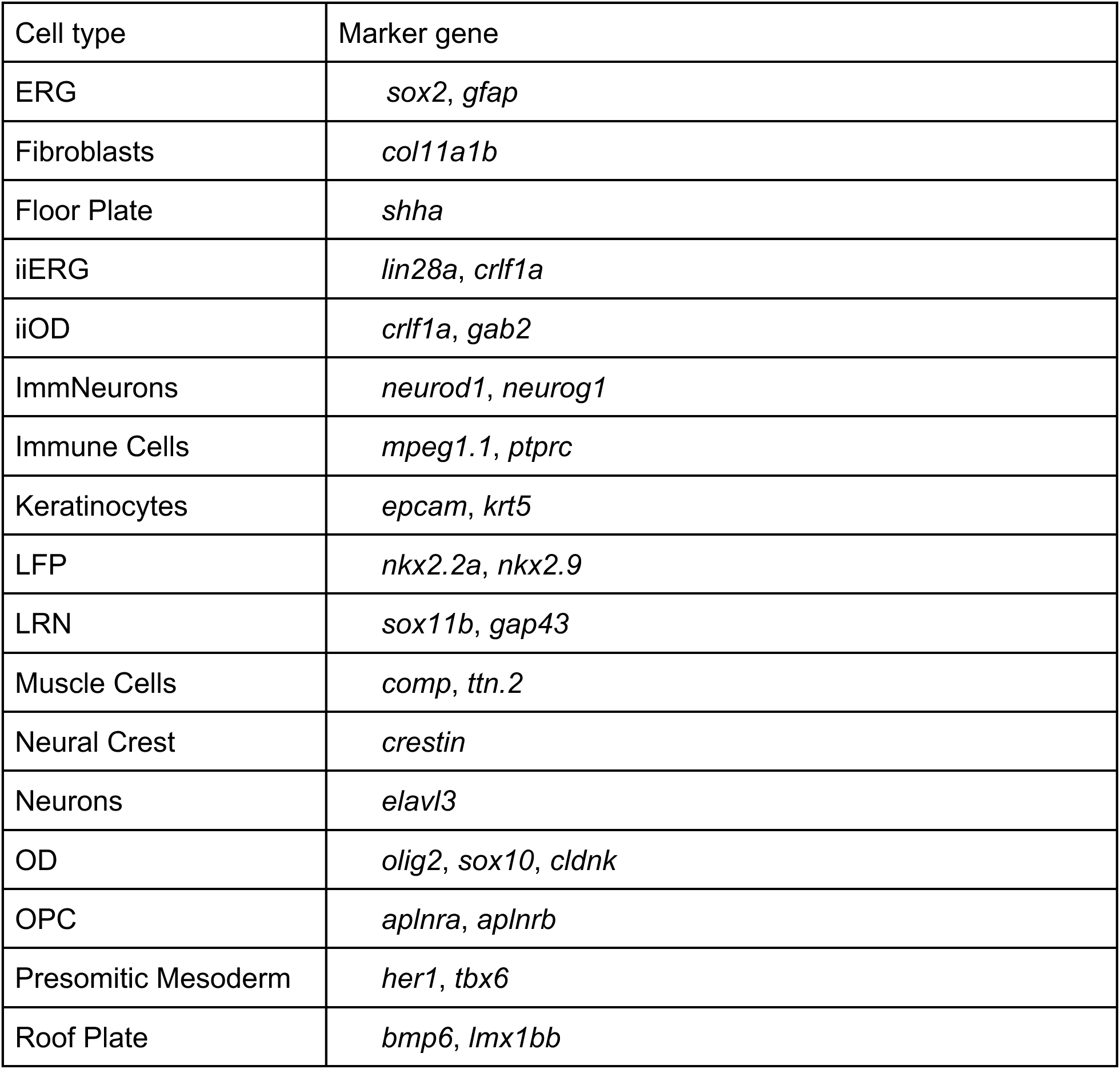
Marker genes used for cell-type annotation. Markers were selected based on previously published studies and the top differentially expressed genes identified for each cluster (*17*, *27*, *99–104*).

| Cell type | Marker gene |
| --- | --- |
| ERG | *sox2, gfap* |
| Fibroblasts | *col11a1b* |
| Floor Plate | *shha* |
| iiERG | *lin28a, crlf1a* |
| iiOD | *crlf1a, gab2* |
| ImmNeurons | *neurod1, neurog1* |
| Immune Cells | *mpeg1.1, ptprc* |
| Keratinocytes | *epcam, krt5* |
| LFP | *nkx2.2a, nkx2.9* |
| LRN | *sox11b, gap43* |
| Muscle Cells | *comp, ttn.2* |
| Neural Crest | *crestin* |
| Neurons | *elavl3* |
| OD | *olig2, sox10, cldnk* |
| OPC | *aplnra, aplnrb* |
| Presomitic Mesoderm | *her1, tbx6* |
| Roof Plate | *bmp6, lmx1bb* |

**Supplementary Figure 1.**
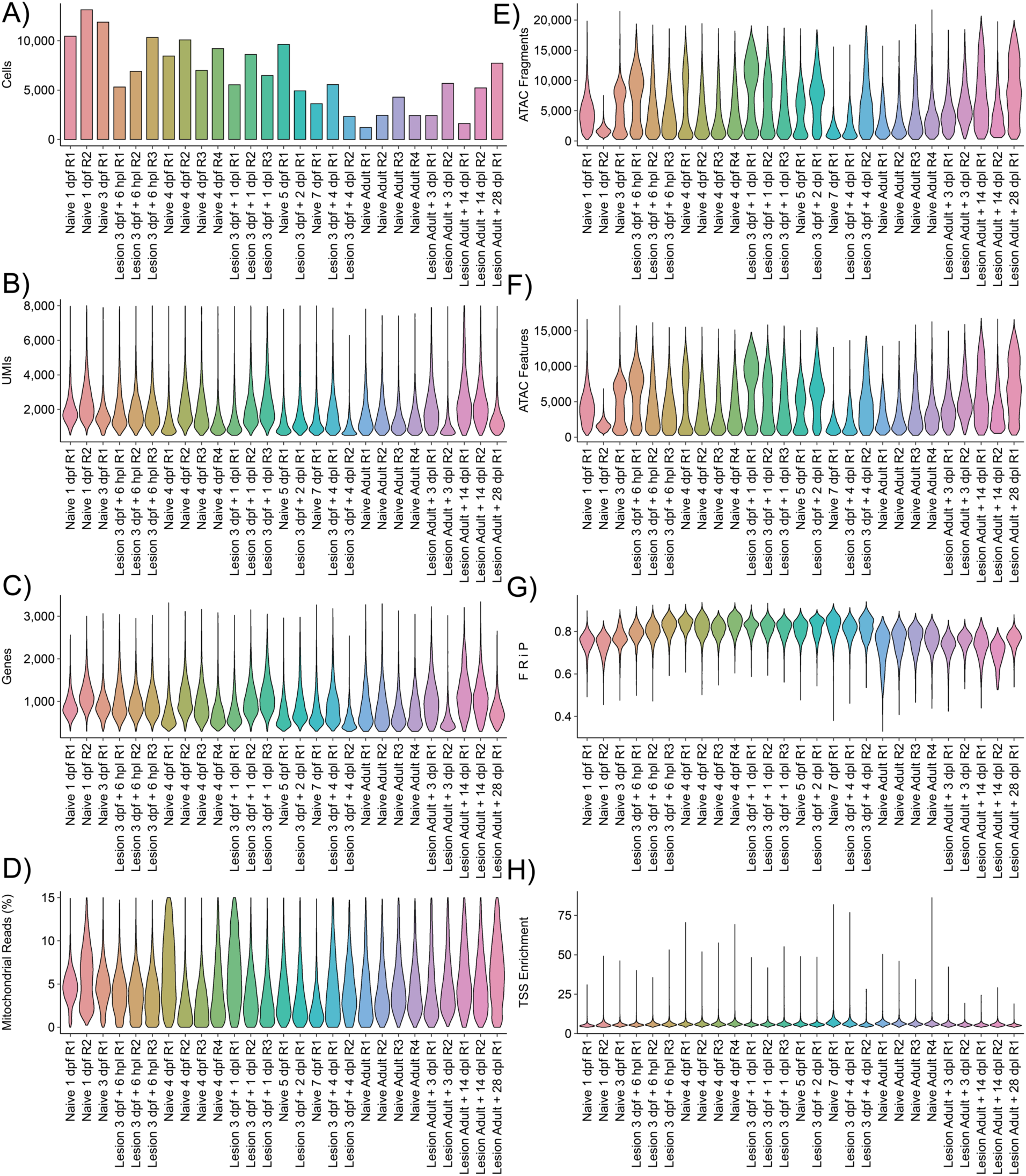
Quality control metrics for the multiome samples. (**A**) Number of cells. (**B**) Number of UMIs, log10 scaled. (**C**) Number of genes, log10 scaled. (**D**) Percentage of reads mapped to the mitochondrial genome. (**E**) Number of ATAC fragments, log10 scaled. (**F**) Number of unique peaks, log10 scaled. (**G**) Number of fragments in peaks. (**H**) TSS enrichment. UMI – unique molecular identifier; dpf – days post fertilization, dpl – days post lesion, hpl – hours post lesion.

**Supplementary Figure 2.**
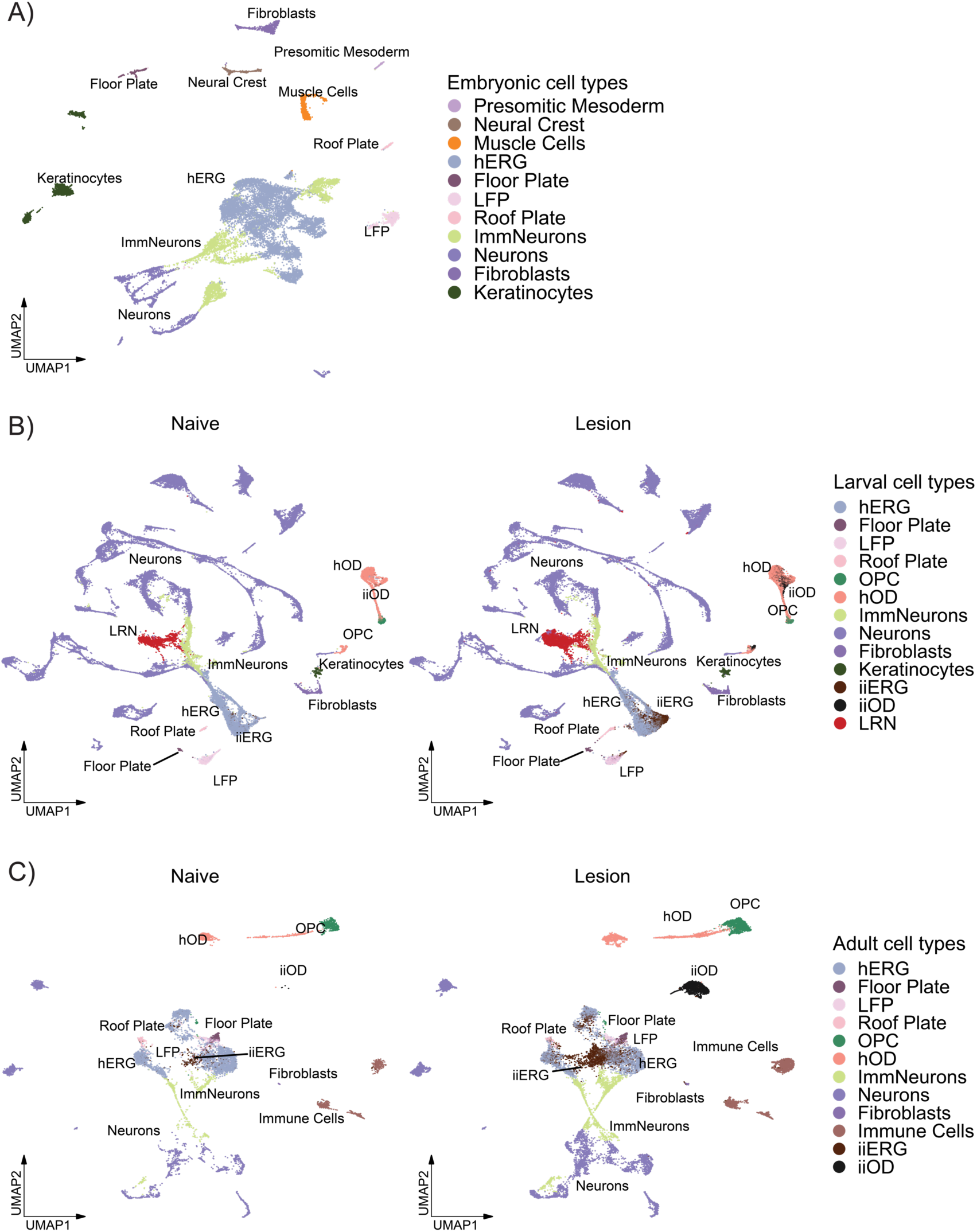
Cell type annotation of each developmental stage. Labels were transferred to the joint atlas. (**A**) UMAP embeddings of 23596 embryonic cells. (**B**) UMAP embeddings of 115835 larval cells split by condition. (**C**) UMAP embeddings of 33002 adult cells split by condition. hERG - homeostatic ependymo-radial glia; hOD - homeostatic oligodendrocytes; iiERG - injury-induced ERG; ImmNeurons - immature neurons; iiOD - injury-induced OD; LFP - lateral floor plate; LRN - lesion-reactive neurons; OPC - oligodendrocyte precursor cells.

**Supplementary Figure 3.**
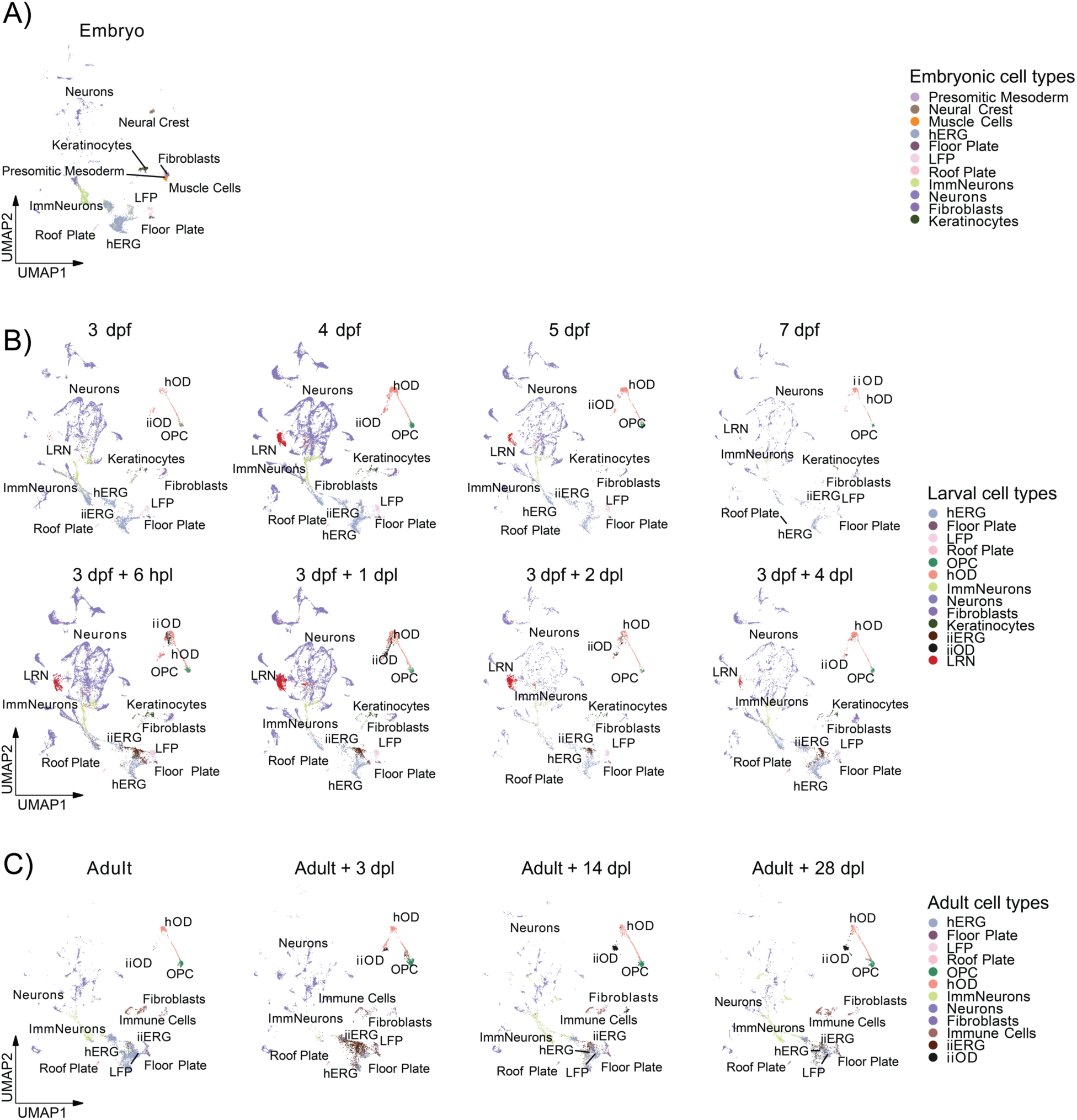
Transferred cell type annotations to the joint atlas. (**A**) UMAP embeddings of embryonic samples. (**B**) UMAP embeddings of larval samples split by time point. (**C**) UMAP embeddings of adult samples split by time point. The top row represents samples from uninjured fish, the bottom row represents samples from injured fish. hERG - homeostatic ependymo-radial glia; hOD - homeostatic oligodendrocytes; iiERG - injury-induced ERG; ImmNeurons - immature neurons; iiOD - injury-induced OD; LFP - lateral floor plate; LRN - lesion-reactive neurons; OPC – oligodendrocyte. dpf – days post fertilization, dpl – days post lesion, hpl – hours post lesion.

**Supplementary Figure 4.**
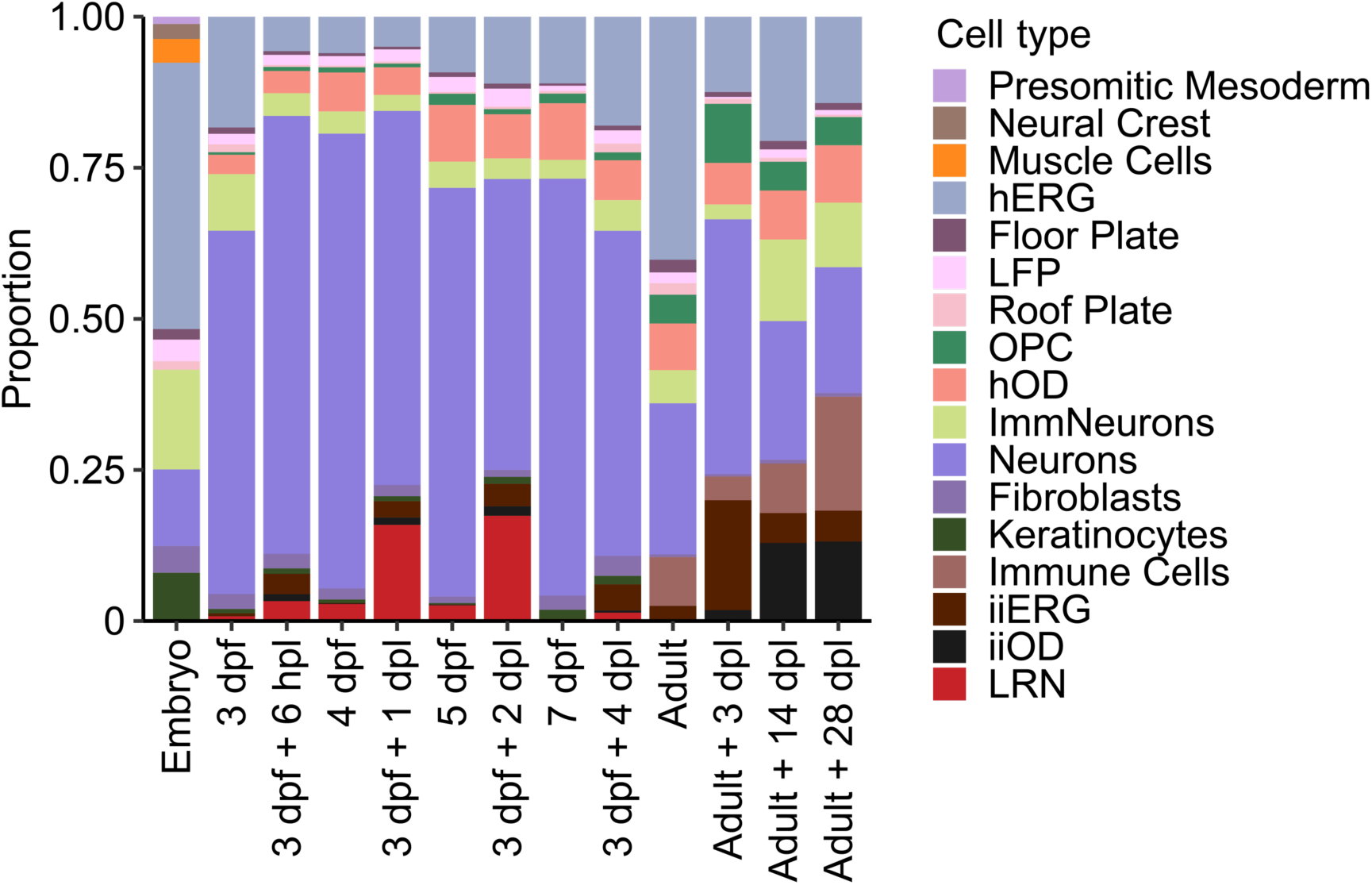
Proportion of cell types in each time point. 3 dpf, 4 dpf, 5 dpf, 7 dpf and Adult correspond to the control samples; 3 dpf + 6 hpl, 3 dpf + 1 dpl, 3 dpf + 2 dpl, 3 dpf + 4 dpl correspond to the spinal cord injury samples. hERG - homeostatic ependymo-radial glia; hOD - homeostatic oligodendrocytes; iiERG - injury-induced ERG; ImmNeurons - immature neurons; iiOD - injury-induced OD; LFP - lateral floor plate; LRN - lesion-reactive neurons; OPC – oligodendrocyte. dpf – days post fertilization, dpl – days post lesion, hpl – hours post lesion.

**Supplementary Figure 5.**
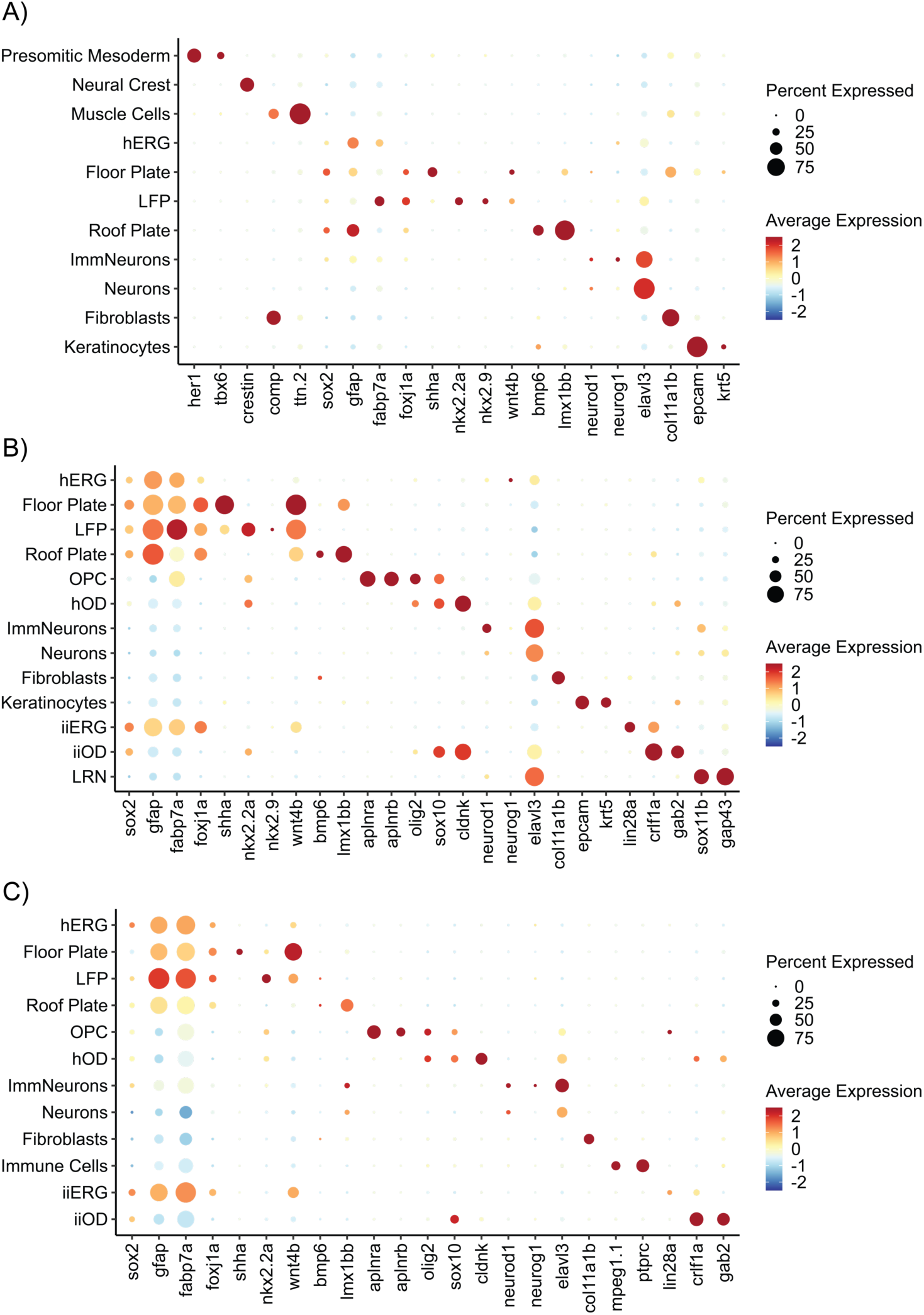
Average scaled expression of marker genes for each identified major cell type. (**A**) Gene expression for embryonic samples. (**B**) Gene expression for larval samples. (**C**) Gene expression for adult samples. hERG - homeostatic ependymo-radial glia; hOD - homeostatic oligodendrocytes; iiERG - injury-induced ERG; ImmNeurons - immature neurons; iiOD - injury-induced OD; LFP - lateral floor plate; LRN - lesion-reactive neurons; OPC – oligodendrocyte. dpf – days post fertilization, dpl – days post lesion, hpl – hours post lesion.

**Supplementary Figure 6.**
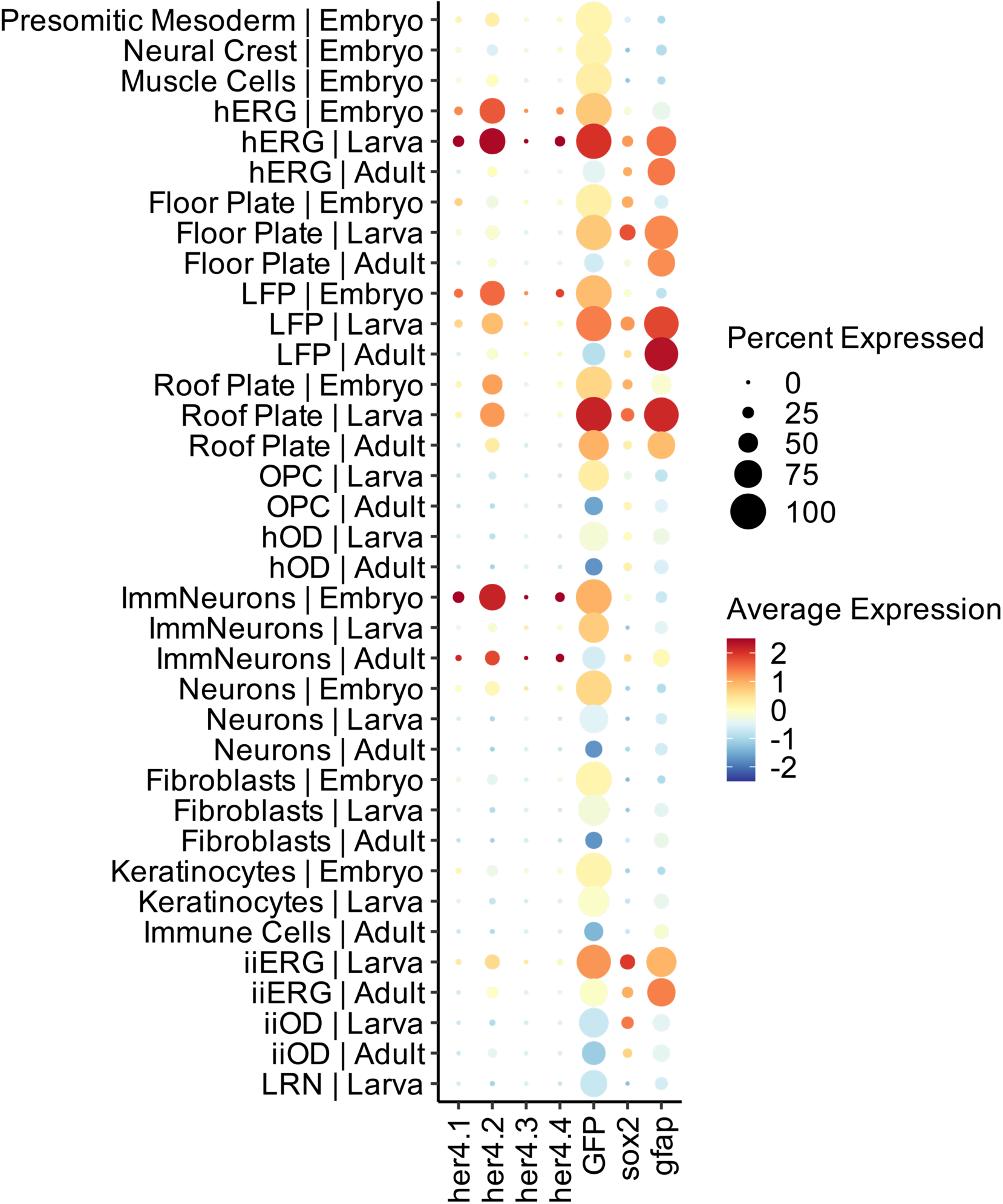
Average scaled expression of her4 genes, GFP and spinal cord progenitor marker genes in each cell type across developmental stages. hERG - homeostatic ependymo-radial glia; hOD - homeostatic oligodendrocytes; iiERG - injury-induced ERG; ImmNeurons - immature neurons; iiOD - injury-induced OD; LFP - lateral floor plate; LRN - lesion-reactive neurons; OPC – oligodendrocyte.

**Supplementary Figure 7.**
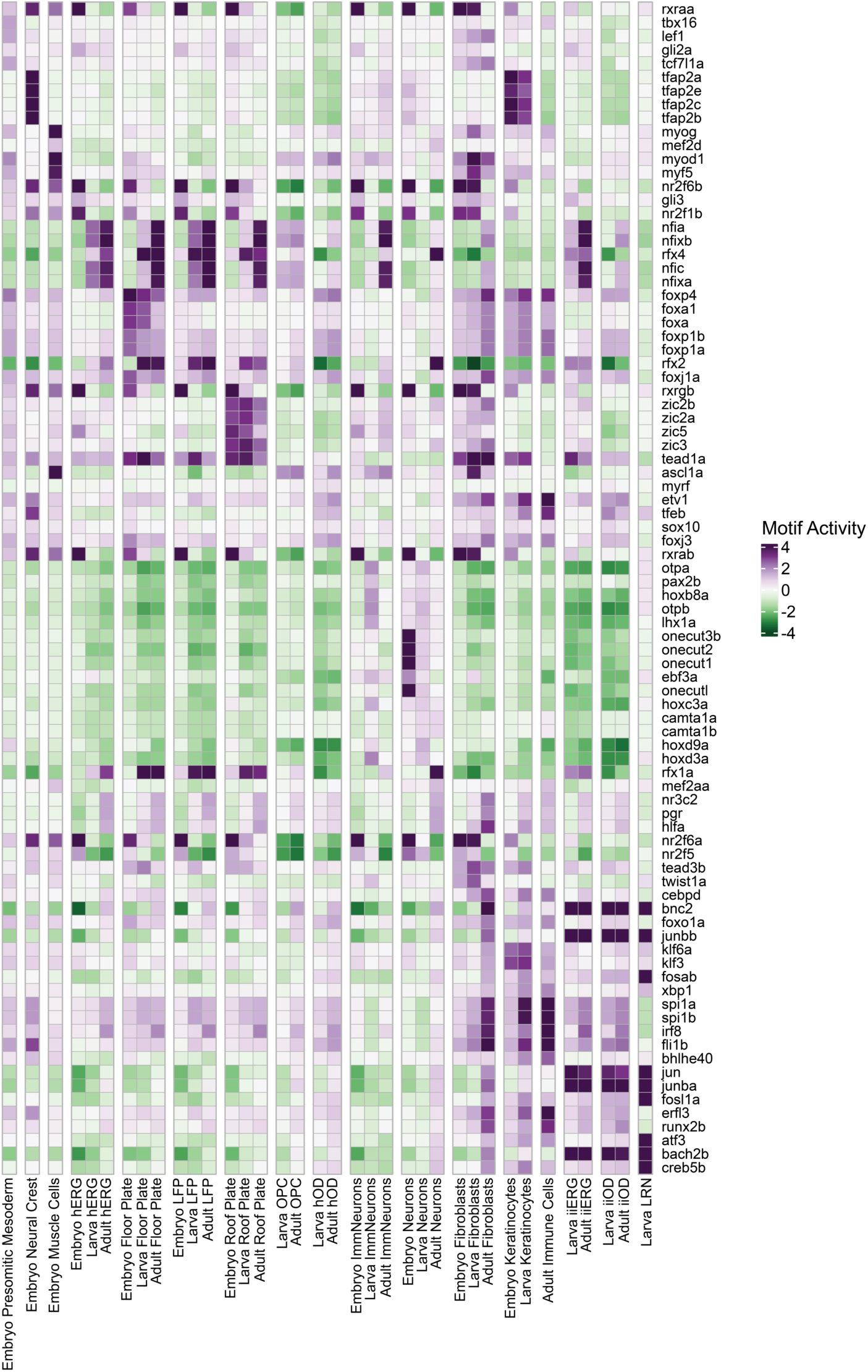
Top five active motifs in major cell types at each developmental stage. The selection was done based on the average motif activity and corresponding TF expression. hERG - homeostatic ependymo-radial glia; hOD - homeostatic oligodendrocytes; iiERG - injury-induced ERG; ImmNeurons - immature neurons; iiOD - injury-induced OD; LFP - lateral floor plate; LRN - lesion-reactive neurons; OPC – oligodendrocyte.

**Supplementary Figure 8.**
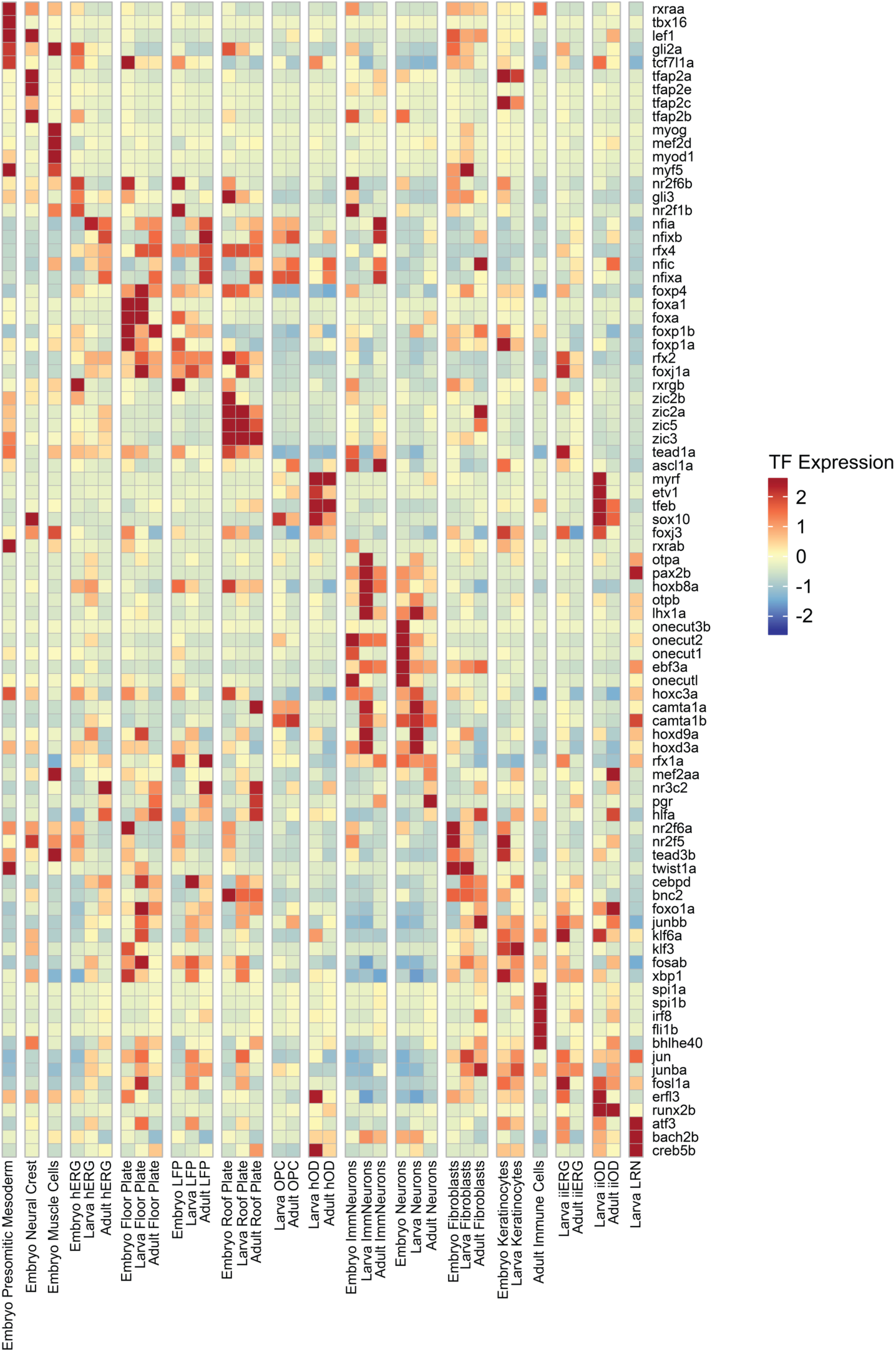
Average scaled expression of selected TFs identified from motif activity analysis. The selection was done based on the average motif activity and corresponding TF expression. hERG - homeostatic ependymo-radial glia; hOD - homeostatic oligodendrocytes; iiERG - injury-induced ERG; ImmNeurons - immature neurons; iiOD - injury-induced OD; LFP - lateral floor plate; LRN - lesion-reactive neurons; OPC – oligodendrocyte.

**Supplementary Figure 9.**
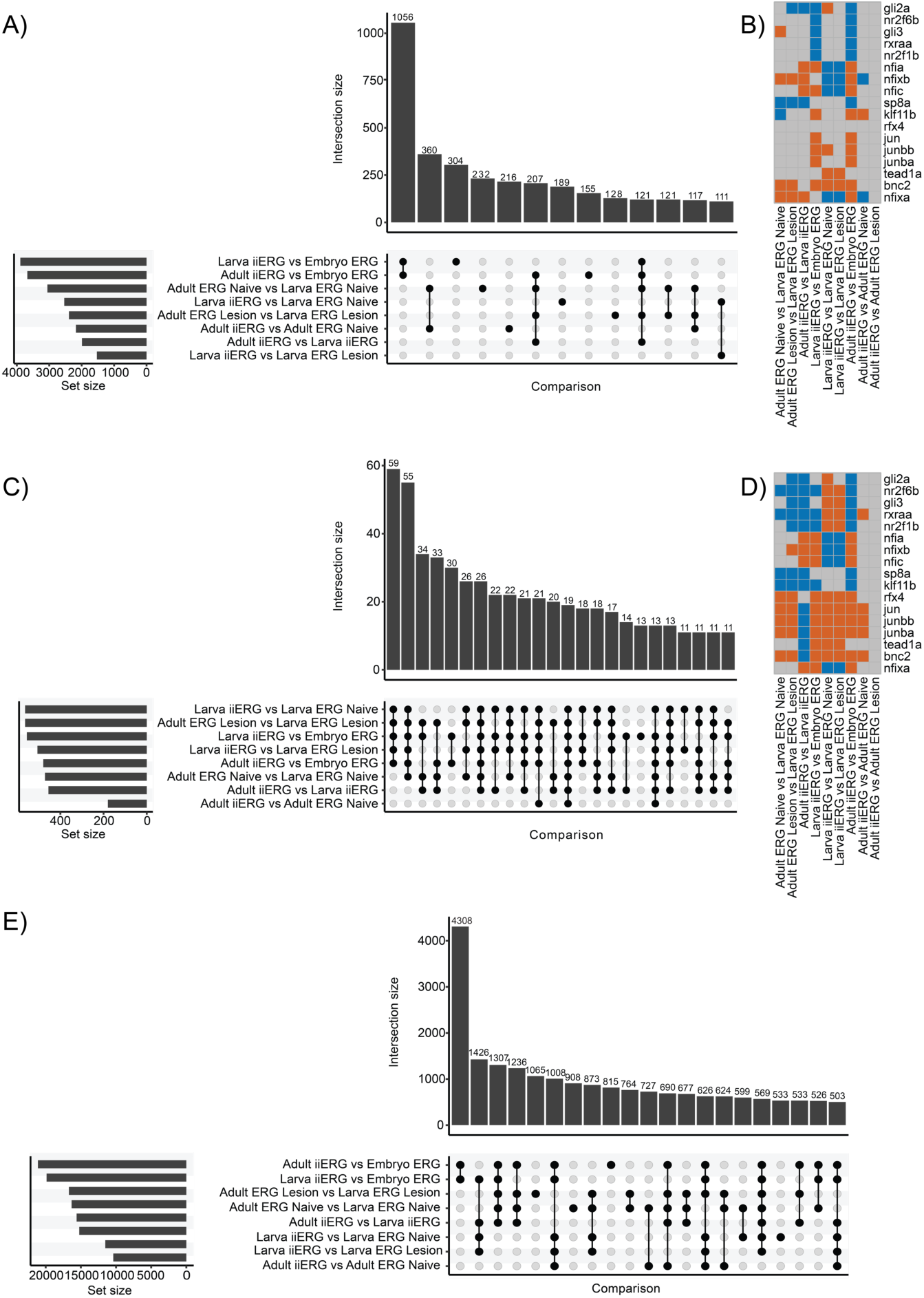
Differential analysis for progenitor cells between developmental stages. Comparisons are annotated as group 1 vs group 2. (**A**) Upset plot of differential motif activity analysis. (**B**) Differential TF motif activity analysis heatmap for top 5 active motifs in each progenitor cell groups; orange color - significantly upregulated in the group 1; blue color - significantly upregulated in the group 2. (**C**) Upset plot of differential gene expression analysis. (**D**) Differential gene expression analysis heatmap for top 5 active corresponding to motifs TFs in each progenitor cell groups; orange color - significantly upregulated in the group 1; blue color - significantly upregulated in the group 2. (**E**) Upset plot of differential accessibility analysis. hERG - homeostatic ependymo-radial glia; iiERG - injury-induced ERG.

**Supplementary Figure 10.**
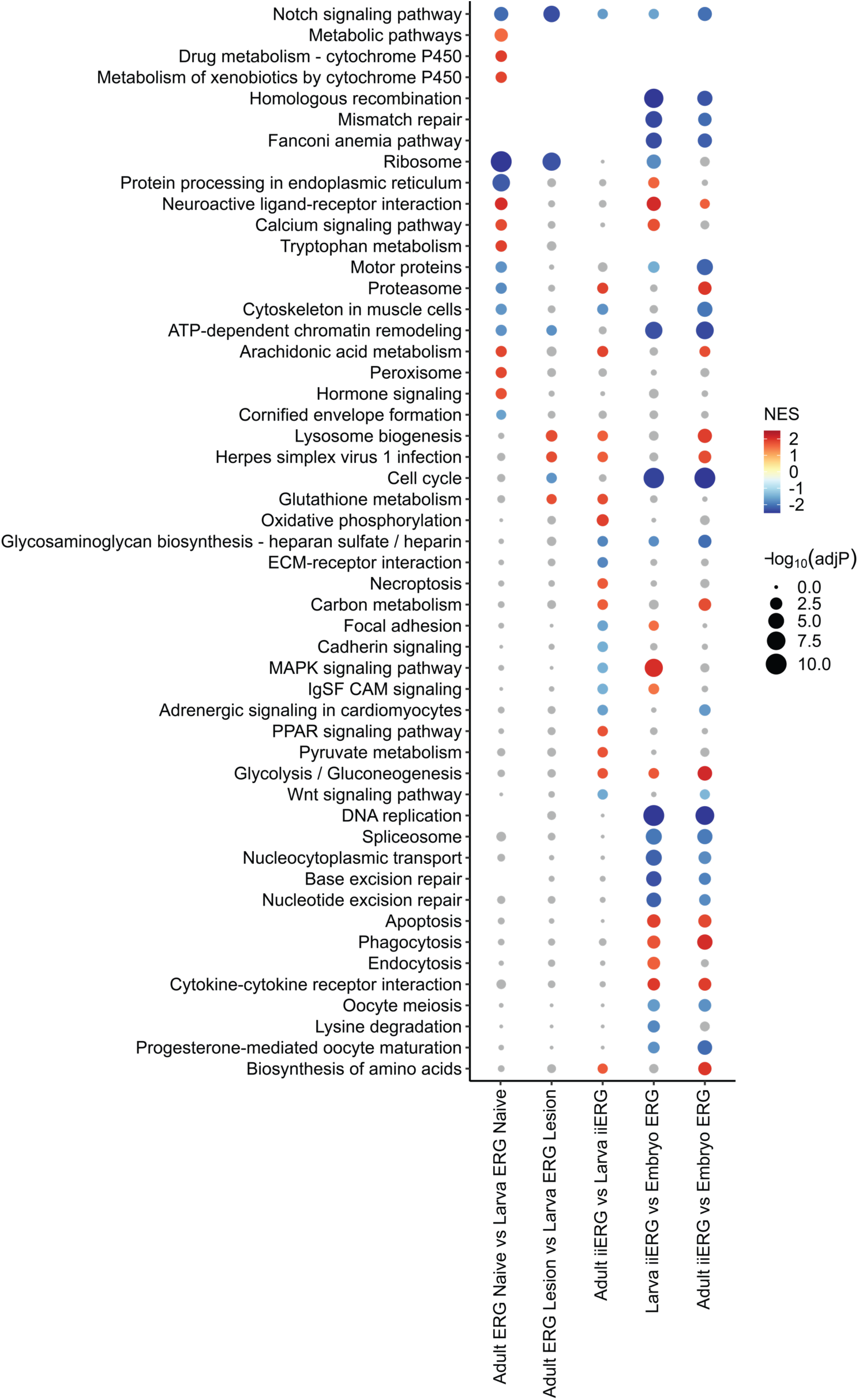
KEGG pathway GSEA analysis on DEGs derived from the developmental stage comparisons across cell groups. hERG - homeostatic ependymo-radial glia; iiERG - injury-induced ERG.

**Supplementary Figure 11.**
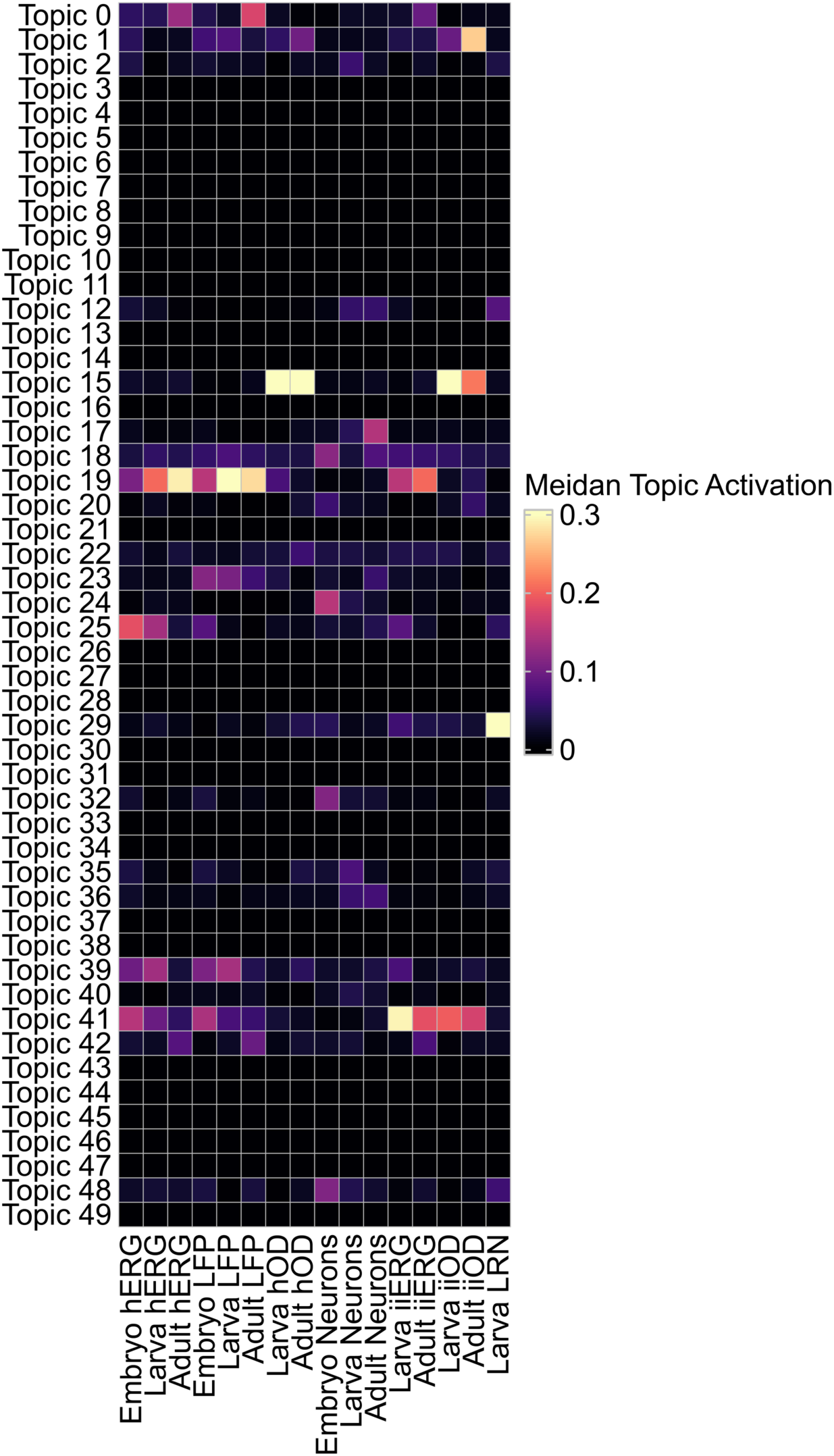
Median topic activity for GRN inference selected cell types across developmental stages. hERG - homeostatic ependymo-radial glia; hOD - homeostatic oligodendrocytes; iiERG - injury-induced ERG; iiOD - injury-induced OD; LFP - lateral floor plate; LRN - lesion-reactive neurons.

**Supplementary Figure 12.**
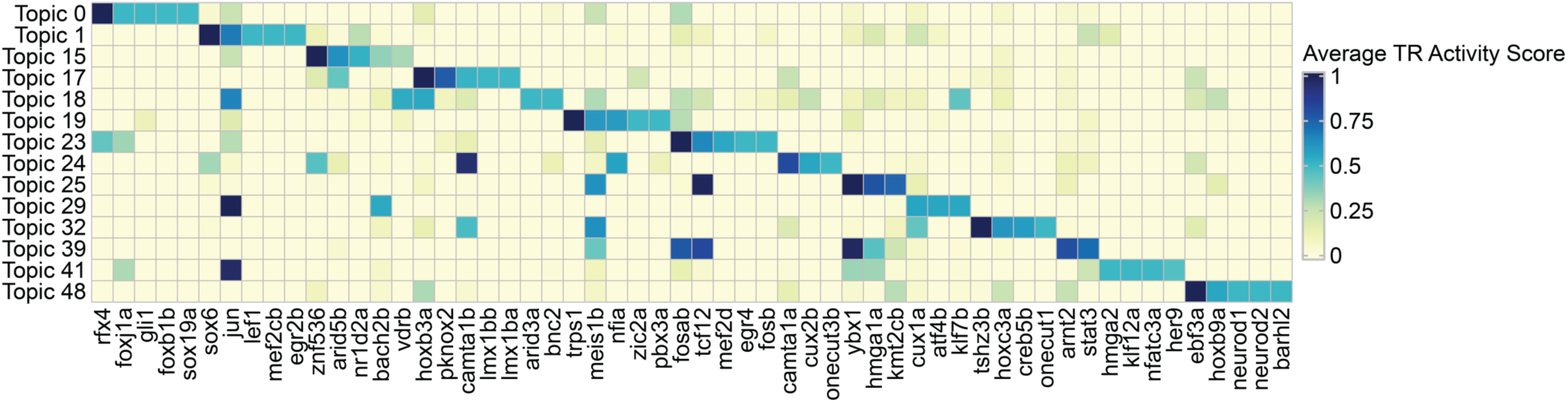
Top 5 TRs for each Topic. Topics that passed activity threshold of 0.1 are shown.

**Supplementary Figure 13.**
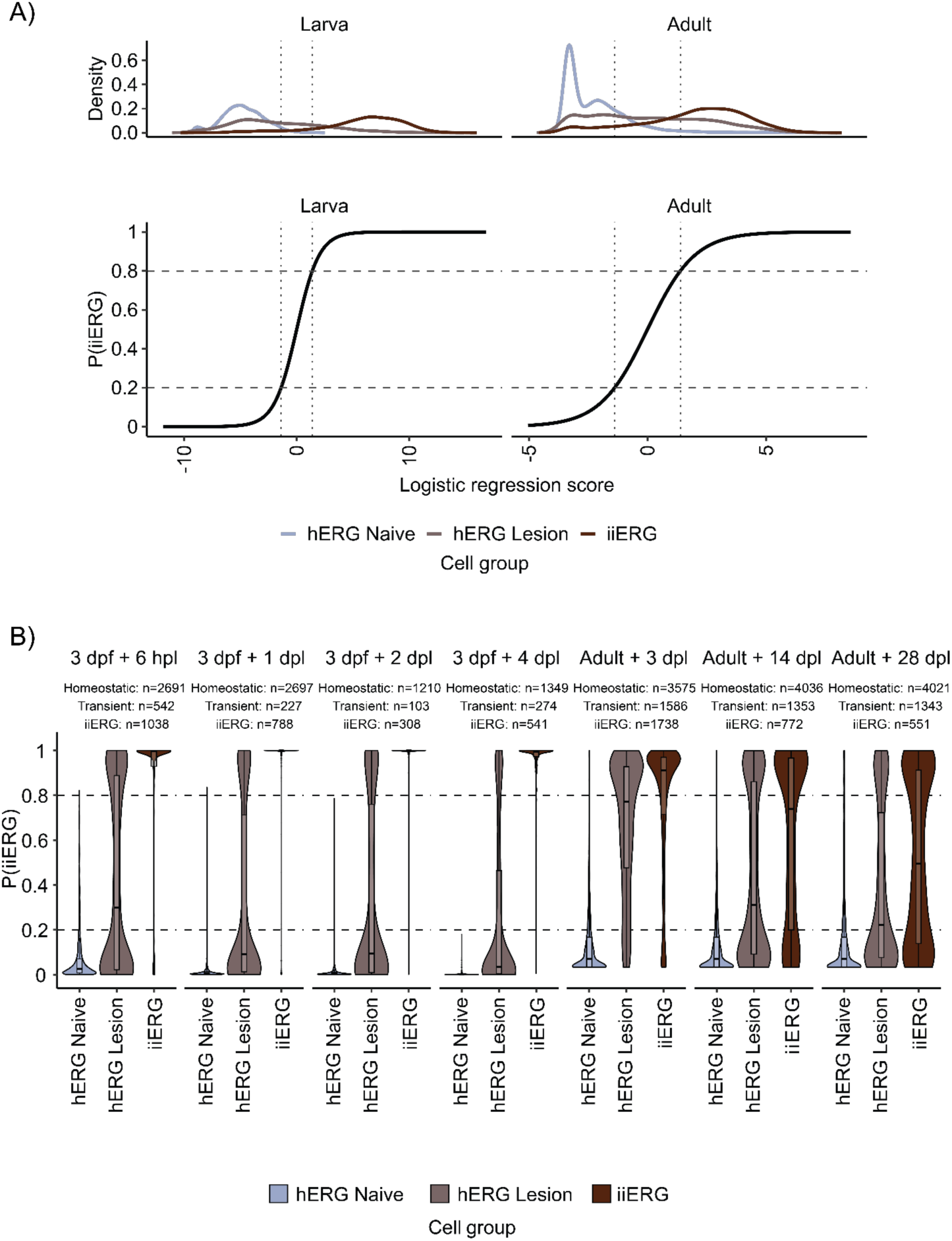
Logistic regression classifies progenitor cells into cell states based on the injury-associated Topic 41. (**A**) Top panel: Distribution of ERG cells along the logistic regression score axis. Bottom panel: Sigmoid curve of the logistic regression classifier. Horizontal lines represent the cut-offs for the cell state classification. (**B**) Classification of ERG cells based on the logistic regression classifier within each time point. To account for the absent matching control time point, ERG cells from 3 dpf and 3 dpf + 6 hpl samples were pooled and cells from Adult Naive samples were added to the corresponding adult Lesion samples. hERG – homeostatic ependymo-radial glia; iiERG - injury-induced ERG; dpf – days post fertilization, dpl – days post lesion, hpl – hours post lesion.

**Supplementary Figure 14.**
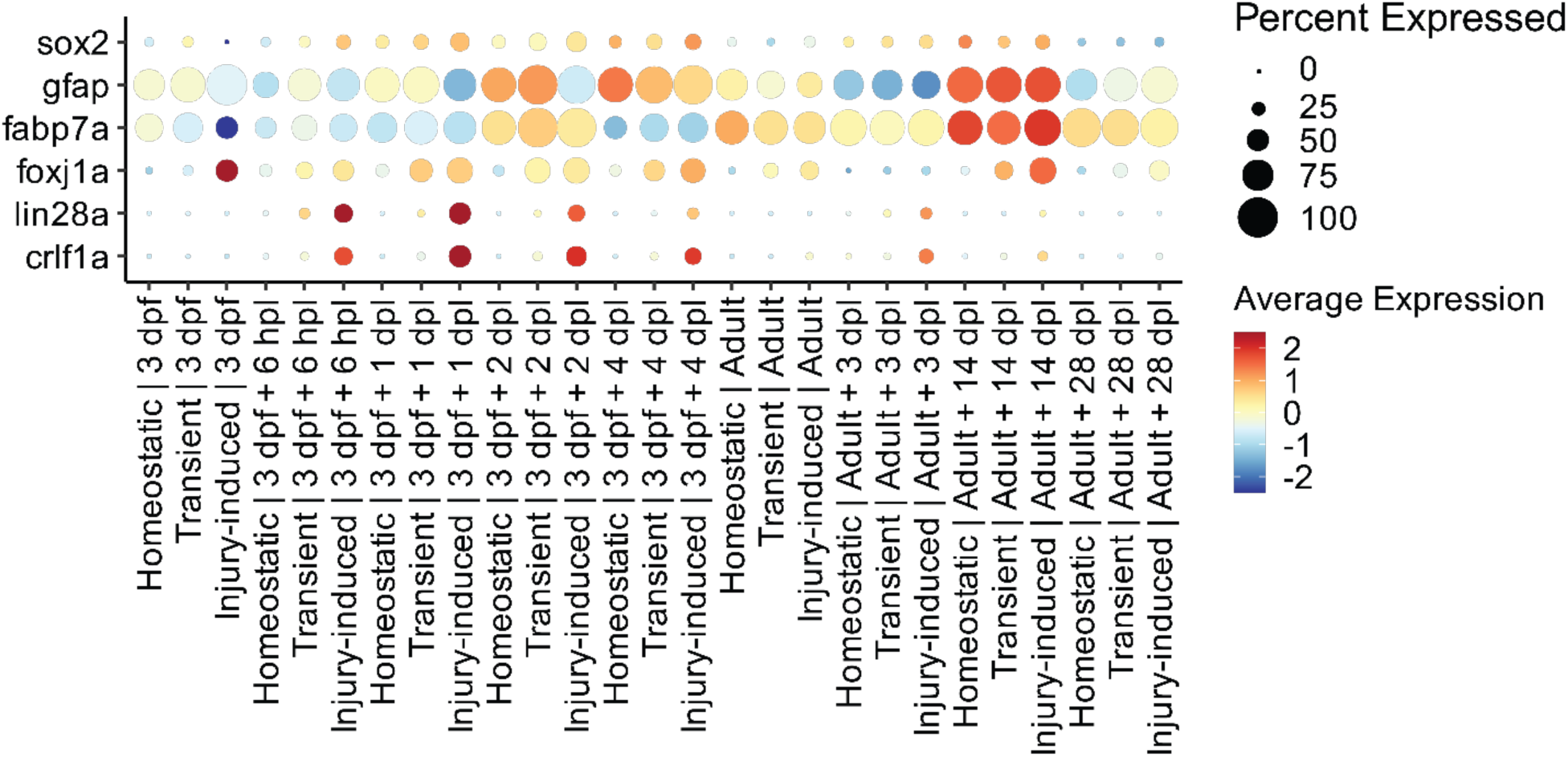
Average scaled expression of hERG and iiERG markers for each cell state across time points. hERG - homeostatic ependymo-radial glia; iiERG - injury-induced ERG; dpf – days post fertilization, dpl – days post lesion, hpl – hours post lesion.

**Supplementary Figure 15.**
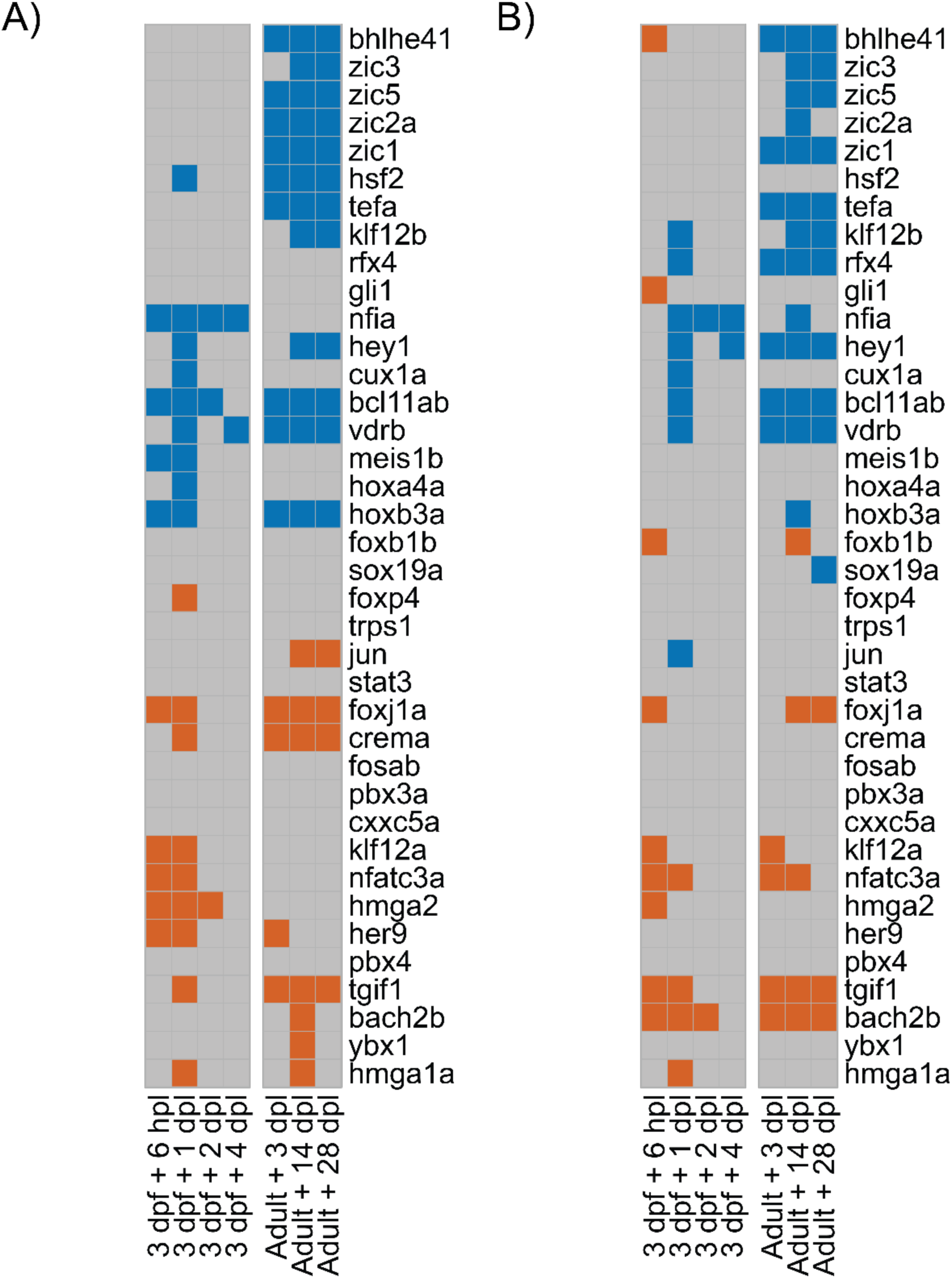
Differential analysis between injury-induced and homeostatic cell states for progenitor cells. (**A**) Differential gene expression analysis heatmap of TRs; orange color - significantly upregulated in the injury-induced cell state; blue color - significantly downregulated in the injury-induced cell state. (**B**) Differential gene and promoter (2kb upstream of the TSS) accessibility analysis heatmap of TRs; orange color - significantly upregulated in the injury-induced cell state; blue color - significantly downregulated in the injury-induced cell state. dpf – days post fertilization, dpl – days post lesion, hpl – hours post lesion.

**Supplementary Figure 16.**
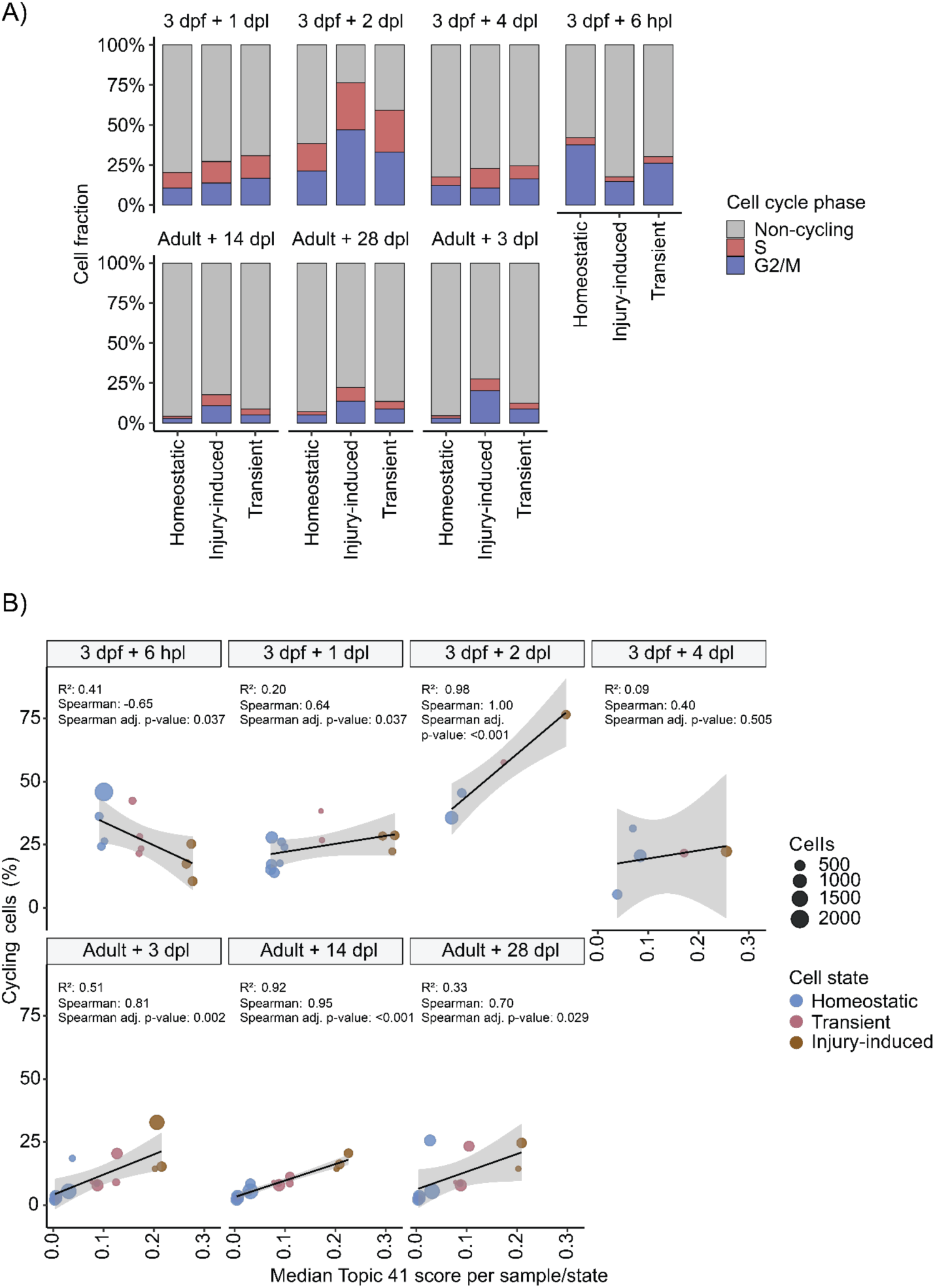
Linear model shows the increase in the percentage of cycling cells in adult samples along the injury-associated Topic 41. To account for the absent matching control time point, ERG cells from 3 dpf and 3 dpf + 6 hpl samples were pooled and cells from Adult Naive samples were added to the corresponding adult Lesion samples. (**A**) Percentage of non-cycling cells and cells in the S and G2/M phase for each cell state at each time point. (**B**) Modelling of percentage of cycling cells at each time point. Each dot represents the sample per time point per cell state combination. ERG – ependymo-radial glia; dpf – days post fertilization, dpl – days post lesion, hpl – hours post lesion.

